# A Multimodal Graph Learning Framework for Versatile Spatial Transcriptomics Analysis with SpatialModal

**DOI:** 10.1101/2025.05.11.653070

**Authors:** Xingyi Li, Dongmin Zhao, Xiangting Jia, Gaoyuan Du, Jialuo Xu, Yang Qi, Yiqi Chen, Yingfu Wu, Junnan Zhu, Feng Wei, Min Li, Xuequn Shang

**Affiliations:** School of Computer Science, Northwestern Polytechnical University, Shannxi, China; State Key Laboratory of Multimodal Artificial Intelligence Systems, Institute of Automation, CAS, Beijing, China; Department of Hepatobiliary and Pancreatic Surgery, General Surgery Center, First Hospital of Jilin University, Jilin, China; School of Computer Science, Central South University, Changsha, Hunan, China

## Abstract

The development of spatial transcriptomics(ST) technologies enables the exploration of tissue structure and cellular function within a spatial context. However, mainstream analytical methods primarily focus on the gene expression modality itself and struggle to fully leverage auxiliary information from histological images, which limits precise dissection of complex biological structures. To address this, we propose SpatialModal, a multimodal graph learning framework that effectively integrates gene expression data and histological images from spatial transcriptomics to construct unified spatial representations. SpatialModal effectively captures synergistic interactions between modalities by jointly modeling intra- and inter-modal complementary features while incorporating spatial adjacency information. Furthermore, it employs a dual contrastive learning strategy to enhance the discriminative power of representations, thereby enabling efficient and robust analysis of tissue structures. We validate the effectiveness of this approach on multiple public datasets covering diverse tissue types, species, and resolutions. Experiments demonstrate that SpatialModal exhibits significant advantages in downstream tasks, including spatial domain identification, gene expression reconstruction, and pseudotime inference, and accurately elucidates the hierarchical structure of the mouse cortex as well as the metabolic-immune dynamic boundaries in the breast cancer microenvironment. Additionally, SpatialModal shows exceptional robustness in single-cell resolution and multi-slice integration tasks, providing an efficient and broadly applicable analytical tool for spatial transcriptomics research.

## Introduction

In multicellular organisms, cells form functional units through highly organized spatial arrangements, collectively maintaining tissue homeostasis and executing complex physiological functions. Dissecting this spatial regulatory mechanism holds significant importance for understanding biological processes such as organ development, immune responses, and tumor microenvironments^1^. Recent breakthroughs in ST^2–4^ technologies enable the acquisition of gene expression data while preserving spatial location information, offering new perspectives for revealing the spatial dynamics of tissue structure and function^5,6^.

Despite this progress, existing analytical methods still face challenges in accurately delineating spatial tissue architecture. For instance, BayesSpace^7^ employs a Bayesian statistical model for fine-grained spatial domain segmentation; GraphST^8^ leverages graph-based self-supervised contrastive learning to enhance expression representations; STAGATE^9^ utilizes a graph attention autoencoder to capture spatial dependencies; STAligner^10^ focuses on batch effect correction across tissue slices. While these methods have demonstrated effectiveness in their respective tasks, they primarily rely on gene expression data for modeling. This reliance limits their capacity to distinguish regions with similar expression profiles but distinct morphological structures, thereby restricting their utility in complex tissue analysis.

Notably, ST experiments typically include high-resolution histological images, particularly hematoxylin and eosin (H&E) stained images. These images provide rich contextual information regarding cellular morphology, extracellular matrix distribution, and vascular structures, offering a potential complement to expression data. Integrating image and expression modalities has thus emerged as a promising direction to improve spatial analysis. Such multimodal fusion not only enhances the resolution of spatial clustering but also strengthens the biological interpretability of functional region annotation, cell type identification, and disease microenvironment analysis. An early representative method, SpaGCN, utilizes RGB values of spatial points in images to assist in calculating edge weights in graph structures. Subsequently, most methods shift toward modality integration by enhancing node features, commonly employing pretrained deep learning models to extract image features. Based on the approach to image feature integration, existing methods broadly fall into two categories: one category uses image features to compute weights for neighboring gene expression, subsequently enhancing gene expression through interpolation, with representative methods including stLearn^11^, DeepST^12^, and ResST^13^. Additionally, STAIG^14^ adopts a graph contrastive learning framework, using image features to calculate edge deletion probabilities across multiple views, indirectly optimizing the expression graph structure. These methods rely on image feature spaces to assist in adjusting gene expression, but image information does not directly contribute to node representation learning. The other category directly incorporates image features into node features, fusing them with gene expression data, as exemplified by EfNST^15^. Although these methods achieve modality-level joint modeling, the fusion process depends on fixed parameter settings, lacking adaptive modeling capabilities for modality differences and local heterogeneity, resulting in limited robustness.

Here, we introduce SpatialModal, a spatial analysis framework with adaptive cross-modal fusion capabilities. Built upon an image-guided augmentation strategy, SpatialModal employs a hierarchical encoding scheme to extract both cross-modality shared information and modality-specific information. Based on these embeddings, we introduce an adaptive fusion mechanism to integrate multimodal features in the latent space, balancing the weights and complementarity of expression and morphological information. Furthermore, we introduce a dual contrastive learning approach that incorporates multi-view and pre-clustering strategies to construct more consistent and discriminative spatial representations. We extensively validate SpatialModal on ST datasets spanning various tissue types, species, and resolutions. Experimental results show that SpatialModal significantly outperforms existing methods in multiple downstream tasks, demonstrating superior performance across platforms and resolutions, and providing an efficient, versatile, and biologically interpretable analysis framework for ST research.

## Results

### Method overview

SpatialModal is a multimodal ST analysis framework based on graph neural networks, designed to effectively integrate gene expression data with histopathological image features. This method treats spatial points as basic units, dividing corresponding tissue slices into multiple image patches and utilizing pretrained neural networks to extract their image features. Simultaneously, it constructs a K-nearest neighbor graph based on spatial coordinates to form spatial adjacency relationships, inputting both gene expression features and image features into the model to establish the foundation for integrated analysis. In terms of representation learning, SpatialModal adopts a hierarchical encoding strategy to comprehensively extract modality-specific and shared information. Specifically, the model first employs two independent graph convolutional networks (GCNs) to separately encode gene expression and image modalities, obtaining their low-dimensional embedding representations. Subsequently, it feeds both modality embeddings into a shared GCN to learn cross-modal joint representations. Finally, it uses a graph variational autoencoder (VGAE) to deeply integrate these low-dimensional representations, enhancing the expressive power and spatial consistency of the fused representations. To further improve the discriminability of gene expression representations, SpatialModal incorporates a graph-based contrastive learning module during the encoding process. This module takes gene embeddings and the spatial adjacency graph as inputs, generats two augmented views by randomly dropping edges and features, and uses a shared GCN with a projection head to produce contrastive embeddings, guiding the model to learn structure-sensitive representations through contrastive loss. Additionally, to mitigate long-range noise interference, the model introduces a pre-clustering operation on low-dimensional embeddings during training, providing a more stable representation foundation for downstream tasks such as spatial domain identification. After modeling the latent space structure, SpatialModal also designs a gene expression reconstruction module based on spatial coordinate mapping. This module takes learned latent representations and spatial coordinates as inputs, outputting a reconstructed gene expression matrix, and establishes the mapping between encoder and decoder through two multilayer perceptrons (MLPs) with identical layer structures. Notably, when the input consists of multiple homologous tissue slices with spatial correspondence, SpatialModal leverages original and newly generated spatial coordinates to reconstruct higher-resolution tissue gene expression profiles in three-dimensional space, further expanding the models potential for multi-slice joint analysis.

### Human DLPFC Dataset Analysis with 10x Visium

To evaluate the performance of SpatialModal in spatial domain identification, we first benchmark it against seven state-ofthe-art methodsincluding DeepST, GraphST, MuCoST^16^, conST^17^, SEDR^18^, SpaGCN, and stLearnon the 10x Visium^19,20^ DLPFC dataset^21^. Quantitative comparisons using four metrics-Adjusted Rand Index (ARI)^22^, Normalized Mutual Information (NMI)^23^, Homogeneity, and Purity Score^24^)-demonstrate SpatialModal’s superior performance across all metrics (Fig. 2b, Supplementary Fig. 2B, Supplementary Tables S1–S3). For slice 151672, SpatialModal achieves the highest ARI of 0.80, substantially outperforming conST (0.65) and SEDR (0.62). We further validate SpatialModal’s effectiveness by visualizing spatial clustering results across 12 DLPFC slices (Fig. 2a, Supplementary Fig. S1), with detailed analysis of three slices (151507, 151672, and 151674) derived from distinct individuals exhibiting notable histological heterogeneity. Results reveal discontinuous spatial domains in SpaGCN and stLearn outputs, along with regional omissions in SEDR, conST, and DeepST. GraphST detects partial domains in slices 151672 and 151674 but generates irregular regions lacking histological coherence. MuCoST shows improved performance yet remains inferior to SpatialModal. Notably, SpatialModal not only resolves clear cortical laminar structures but also precisely identifies Layer 1 and adjacent Layer 2 in slice 151507, demonstrating exceptional spatial clustering and hierarchical identification capabilities.

**Figure 1.**
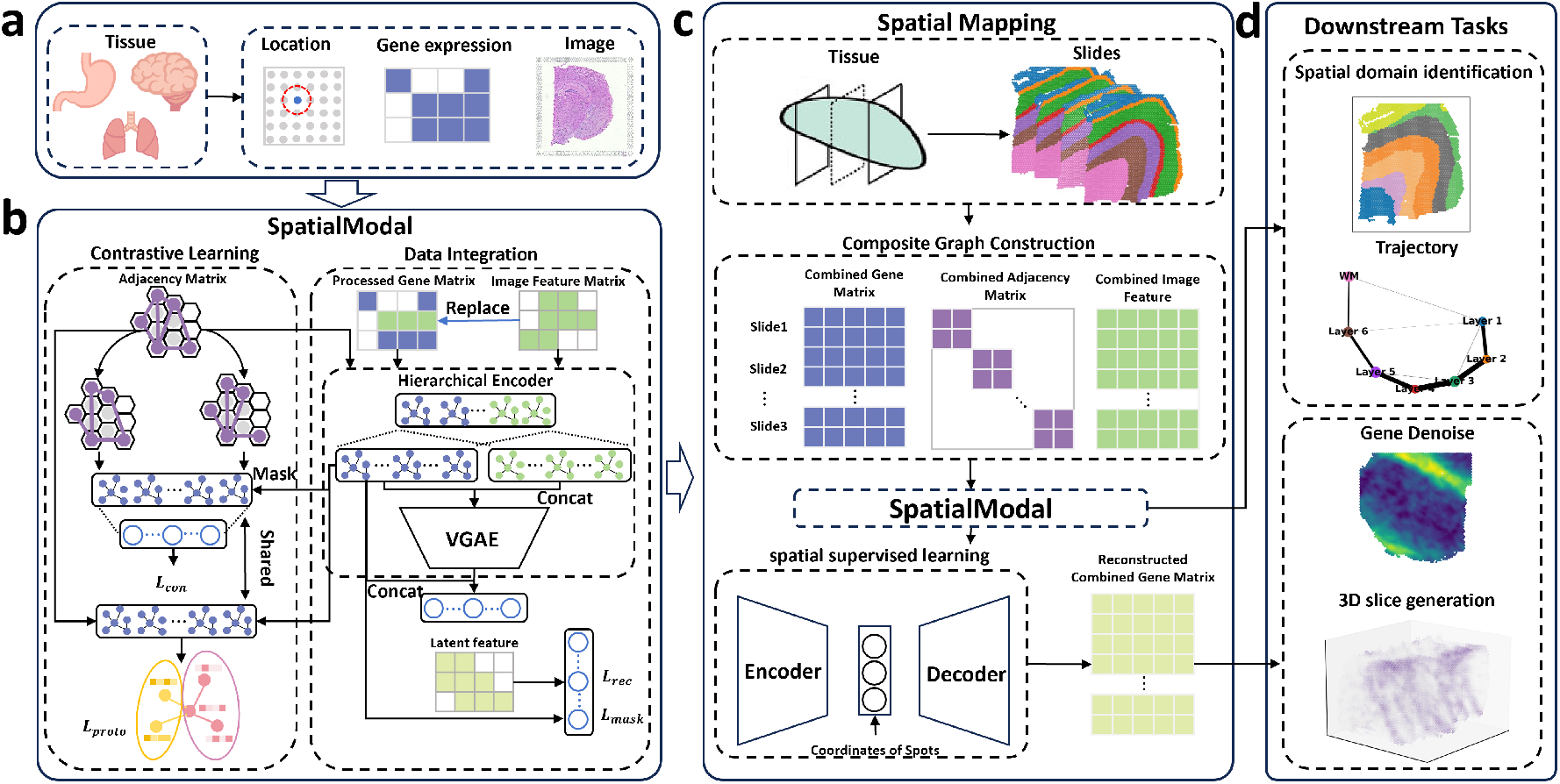
Overview of SpatialModal. a. SpatialModal takes gene expression data, spatial coordinates, and histopathological images from ST as inputs. It segments images around each spatial point and extracts image features using a pretrained ResNet152 model. Simultaneously, it constructs an adjacency matrix based on the coordinates of spatial points. b. It randomly replaces a portion of gene expression vectors with image feature vectors, generating a new matrix that, together with image features, feeds into a shared feature fusion encoder. This encoder consists of a shared graph convolutional network (GCN), two independent GCNs, and a graph variational autoencoder (VGAE). Topology-aware contrastive learning constructs a graph using a low-dimensional gene expression matrix as nodes and the original adjacency matrix as edges, generating augmented graphs by randomly dropping edges and features, and performs pre-clustering on gene expression data. c. In the gene reconstruction phase, it employs a spatial mapping approach. Latent representations undergo processing through a multilayer perceptron (MLP) encoder and decoder. For single-slice training, it generates two-dimensional intermediate vectors and computes spatial loss using spatial coordinates (*x, y*). For multi-slice training, it vertically concatenates gene expression and image feature matrices, diagonally merges adjacency matrices, generates three-dimensional intermediate vectors, and computes spatial loss using spatial coordinates (*x, y*) and slice identifiers. d. SpatialModals generated latent representations support spatial domain identification and trajectory inference. Reconstructed gene expression data enables 2D and 3D visualization analysis.

**Figure 2.**
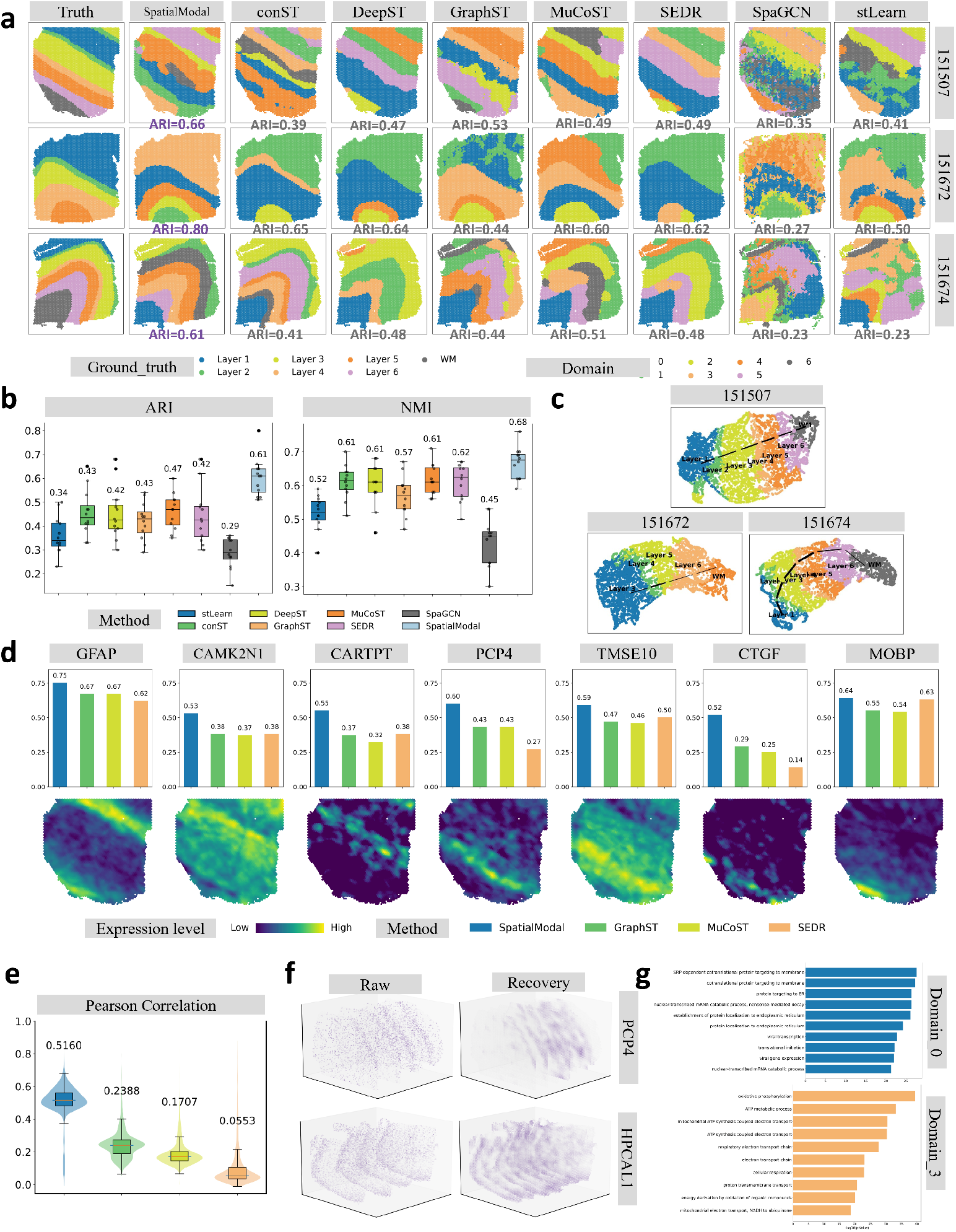
Performance on 10x Visium DLPFC data. a. Ground-truth annotations and spatial clustering results for three representative DLPFC slices across SpatialModal and seven baselines. b. Boxplots comparing ARI and NMI across 12 slices. Medians are indicated by central lines and numerical labels; box limits represent interquartile ranges. c. UMAP visualizations (colored by ground truth) and PAGA-derived developmental trajectories for three slices. d. Spatial patterns of seven layer-specific marker genes reconstructed by SpatialModal in slice 151507. Bar plots show Pearson correlations between reconstructed and raw expression levels for SpatialModal versus three baselines. e. Violin plots comparing global gene expression reconstruction accuracy (Pearson *r*) across methods. f. 3D reconstructions of PCP4 and HPCAL1 expression in slices 151507 and 151674. g. Top 10 GO terms enriched in domains 0 and 3 identified by SpatialModal.

To further validate these advantages, we perform trajectory analysis using UMAP^25^ and PAGA^26^ based on learned latent representations from all methods (Fig. 2c, Supplementary Fig. 2A). SpatialModal uniquely reconstructs a continuous trajectory from Layer 1 through Layer 2 to white matter (WM) in slice 151507, with UMAP visualizations closely matching manual annotations. In slices 151672 and 151674, while SpatialModal, conST, and DeepST generate linear trajectories, the latter two exhibit overlapping boundaries between Layer 4 and adjacent layers, failing to distinguish laminar architecture.

Gene expression recovery analysis of seven known laminar marker genes (Fig. 2d, Supplementary Fig. 3A) confirms SpatialModal’s superior reconstruction fidelity, supported by Pearson correlation coefficients surpassing all baselines (Fig. 2e). The 3D reconstruction by SpatialModal further highlights enhanced resolution and clarity in gene expression patterns (Fig. 2f), accurately reflecting intra-tissue heterogeneity. Finally, GO enrichment analysis of five spatial domains identified in slice 151672 (Fig. 2g, Supplementary Fig. 3B) reveals biologically meaningful functional partitioning: Domains 2–3 show synaptic transmission-related enrichment, aligning with their roles in working memory and executive function; Domain 1 exhibits energy metabolism signatures, reflecting high neuronal energy demands; Domains 4–5 display protein synthesis/transport terms, underscoring their roles in neuronal homeostasis.

### Human BRCA Dataset Analysis with 10x Visium

To assess the performance of SpatialModal in heterogeneity analysis, we benchmark it against seven existing methods using a 10x Visium human breast cancer dataset (Fig. 3a). Initial results show that when setting domain numbers to 10, most methods (except conST, DeepST, and SEDR, which fail to detect normal regions) achieve reasonable proximity to manual annotations. For finer-grained partitioning aligned with ground-truth annotations, we increase domain numbers to 20. Quantitative evaluations demonstrate the superior alignment of SpatialModal with annotated regions, particularly for domains 2 (IDC_4), 5 (IDC_5), 7 (IDC_3), 12 (DCIS/LCIS_4), 17 (DCIS/LCIS_1), and 18 (DCIS/LCIS_5). In performance metrics, SpatialModal attains the highest ARI (0.64) and NMI (0.72), significantly outperforming stLearn and SEDR, while other methods show suboptimal ARI (*<* 0.6) (Fig. 3b). These results highlight SpatialModal’s ability to delineate complex tissue architectures with high precision.

**Figure 3.**
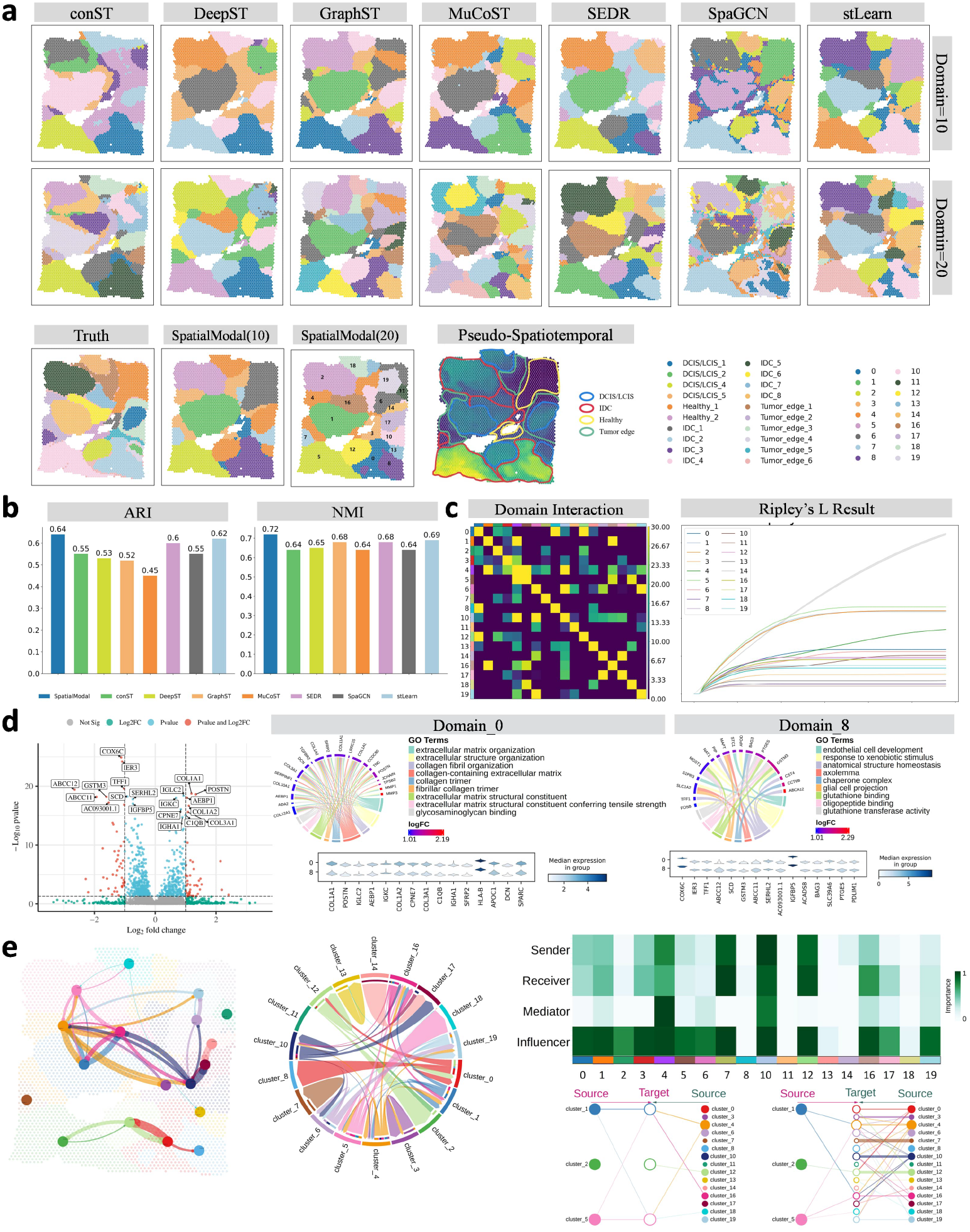
Performance on 10x Visium human breast cancer data. a. Spatial clustering results for seven baselines (*k* = 10, *k* = 20), SpatialModal (*k* = 10, *k* = 20), ground-truth annotations, and pseudotemporal trajectories. b. ARI/NMI comparisons (*k* = 20 clustering). c. Left: Heatmap of inter-domain interaction strengths. Right: Ripley’s *L*-function analysis for spatial domain identification. d. Volcano plot of differentially expressed genes (DEGs) between subregions 0 and 8. Violin plots show expression trends for top DEGs; chord diagram displays enriched GO terms. e. CellChat-based cell communication networks across SpatialModal-identified domains. Heatmaps quantify sender/receiver roles; hierarchical graphs resolve CXCL signaling dynamics.

Using the latent representations from SpatialModal, we construct a pseudospatial-temporal map (pSM)^27^ to resolve interregional heterogeneity (Fig. 3d). domains 2 (IDC_4), 0, and 8 (subdivided from IDC_2) exhibit strong heterogeneity, contrasting with relatively uniform tumor domains. Notably, the subdivision of IDC_2 into domains 0 and 8 by SpatialModal aligns with tendencies observed in GraphST, SpaGCN, stLearn, and SEDR. Quantification of inter-regional interaction strength identifies significant crosstalk between domains 0 and 8 (Fig. 3c), suggesting functional interdependence in the tumor microenvironment. Differential expression analysis identifies 64 genes (*z*-score *>* 3, adjusted *p*-value *<* 10^*−*3^, log_2_FC *>* 1) distinguishing these subregions (Fig. 3d, Supplementary Tables S4–S5). SpatialModal’s reconstructed expression profiles show high concordance with raw data while smoothing noise-corrupted domains (Supplementary Fig. S4). Enrichment analysis reveals domain 0 upregulates genes associated with extracellular matrix remodeling (e.g., *COL1A1, COL1A2, COL3A1, POSTN*) and immune/complement activation (e.g., *IGLC2, IGKC, IGHA1, C1QB*), whereas domain 8 enriches metabolic reprogramming (*GSTM3, SCD, COX6C*) and antioxidative defense genes, consistent with its aggressive phenotypic features.

To uncover spatial patterns in breast cancer progression, Ripley’s *L*^28^-function analysis identifies three prominent clusters: domains 1 (DCIS/LCIS_1), 2 (IDC_4), and 5 (IDC_5) (Fig. 3c). CellChat^29^-based communication analysis (Fig. 3e) reveals heightened CXCL pathway activity (specifically CXCL12-ACKR3 and CXCL12-CXCR4 pairs) in IDC domains (2 and 5) versus DCIS/LCIS (domain 1), correlating with tumor invasiveness and immune modulation. domain 1 acts as a communications hub in the CXCL network, suggesting its regulatory role in early-stage tumorigenesis. Intriguingly, spatial heterogeneity persists within IDC subregions, with domain 5 displaying enhanced signaling activity compared to domain 2, potentially representing a distinct aggressive IDC subtype. These findings align with pseudospatial-temporal map trajectories, further validating SpatialModal’s power in resolving tumor microenvironmental dynamics.

### Mouse Brain Datasets Analysis with 10x Visium

To systematically evaluate the performance of SpatialModal in murine brain structure analysis, we first apply it to a mouse forebrain dataset. SpatialModal achieves superior clustering performance with the highest ARI (0.466) and NMI (0.771) (Fig. 4c). In the hindbrain dataset, SpatialModal resolves finer laminar partitioning in the visual cortex (VIS). Despite manual annotation defining six cortical layers, SpatialModal subdivides VIS into four distinct subregions (domains 11, 14, 4, and 13), demonstrating enhanced spatial resolution compared to methods producing only two (stLearn, conST) or three layers

**Figure 4.**
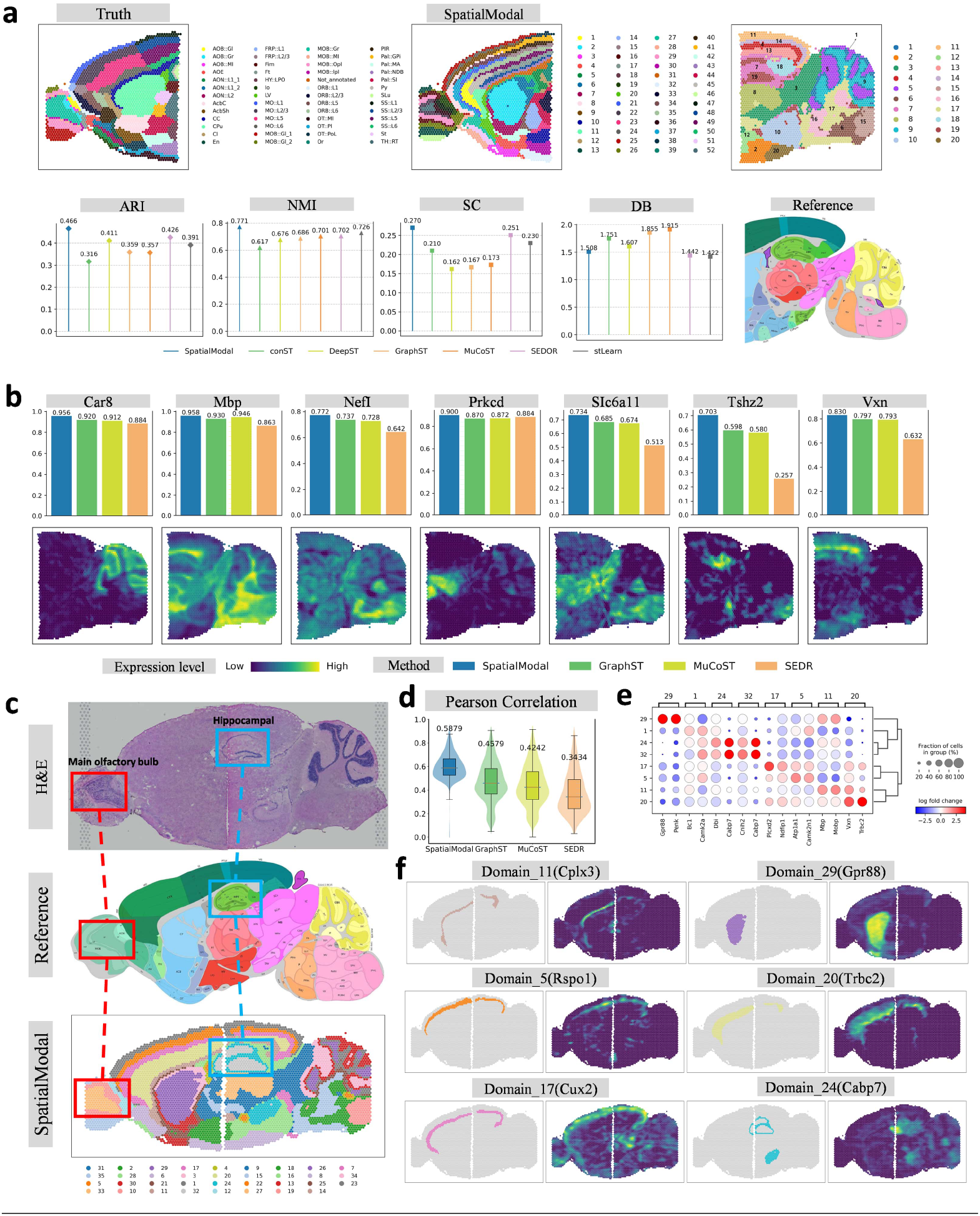
Performance on 10x Visium mouse brain data. a. Ground-truth annotations and SpatialModal clustering for the anterior brain dataset. b. SpatialModal clustering for the posterior brain dataset, alongside Allen Brain Reference Atlas annotations. c. Lollipop plots comparing ARI/NMI (anterior) and Silhouette/Davies-Bouldin scores (posterior). d. Spatial expression patterns of seven posterior brain marker genes. Bar plots show reconstruction accuracy (Pearson *r*). e. H&E-stained anterior/posterior sections, Allen Brain annotations, and SpatialModal-derived domains. f. Violin plots comparing global gene reconstruction performance. g. Bubble plot of differentially expressed genes (DEGs) in cortex and hippocampal regions. h. Spatial gradients of DEG expression aligned with domain boundaries.

(GraphST, MuCoST, SEDR) (Supplementary Fig. S5A). Notably, SpatialModal uniquely identifies the Subiculum and PostSubiculum (domain 18)—a microarchitecture unrecognized by other methods. It further accurately delineates hippocampal domains (domains 19, 7), midbrain (domain 3), thalamus (domain 8), and fiber tracts (domain 17) (Fig. 4b), confirming its precision in structural mapping.

To validate the gene expression reconstruction capability of SpatialModal, we analyze seven spatially clustered domains in the hindbrain dataset and identify key differentially expressed genes (*Car8, Mbp, Nefl, Prkce, Slc6a11, Tshz2*, and *Vxn*). SpatialModal outperforms baseline methods in reconstructing Tshz2 expression patterns (Fig. 4d), explaining its accurate identification of Subiculum/PostSubiculum architecture (domain 18) and showcasing molecular-level resolution.

For multi-slice integration, we jointly train SpatialModal on forebrain and hindbrain datasets. The framework robustly resolves laminar cortical structures, olfactory bulb, and hippocampal formations across slices (Fig. 4e). UMAP visualization of latent spaces reveals minor overlaps only at anatomically continuous tissue interfaces, reflecting precise spatial modeling (Supplementary Fig. S5B). GraphST, MuCoST, and SEDR exhibit similar continuity patterns but inferior discriminative power. SpatialModal achieves the highest Pearson correlation in cross-slice gene expression reconstruction, significantly surpassing benchmarks (Fig. 4f). Differential expression analysis of cross-slice cortical and hippocampal domains (Fig. 4g) confirms that SpatialModal-generated spatial boundaries align with high-expression gradients of marker genes (Fig. 4h), further validating its robustness in molecular pattern recovery.

### SSC and HPA Datasets Analysis with osmFISH and MERFISH

To evaluate the performance of SpatialModal’s on single-cell resolution ST data, we apply it to the osmFISH-generated^30^ mouse somatosensory cortex dataset and compare it with six baseline algorithms. SpatialModal achieves superior spatial domain identification accuracy (ARI = 0.53, NMI = 0.61) (Fig. 5a). Visualization shows high concordance between spatial domains identified by SpatialModal and ground-truth annotations (Fig. 5b). Specifically, SpatialModal precisely partitions Layer 2–3 into subregions 9 (medial) and 3 (lateral), whereas DeepST and SEDR merge them into a single domain, and GraphST, MuCoST, and stLearn fail to resolve laminar structures. SpatialModal also exhibits enhanced accuracy in identifying Layer 4 (domain 10) and Layer 6 (domain 8). Pseudotemporal trajectory inference based on latent representations (Fig. 5b) reveals a developmental path extending from white matter (domains 1 and 7) to Layer 6, aligning with white matter’s role in neuronal migration and cortical development. Notably, the internal capsule caudoputamen (domain 4) displays trajectories linking to adjacent structures, consistent with its early developmental role in sensorimotor signal transmission. UMAP visualization of latent spaces (Fig. 5d, Supplementary Fig. S6A) demonstrates distinct domain separation in SpatialModal, contrasting with mixed representations from conST, MuCoST, and stLearn. Differential expression analysis identifies marker genes (*Plp1, Syt6, Rorb, Cpne5, Slc32a1*) with reconstructed expression patterns closely matching raw data distributions (Fig. 5c, 5d, Supplementary Fig. S6B), confirming SpatialModal’s precision in capturing domain-specific molecular signatures.

**Figure 5.**
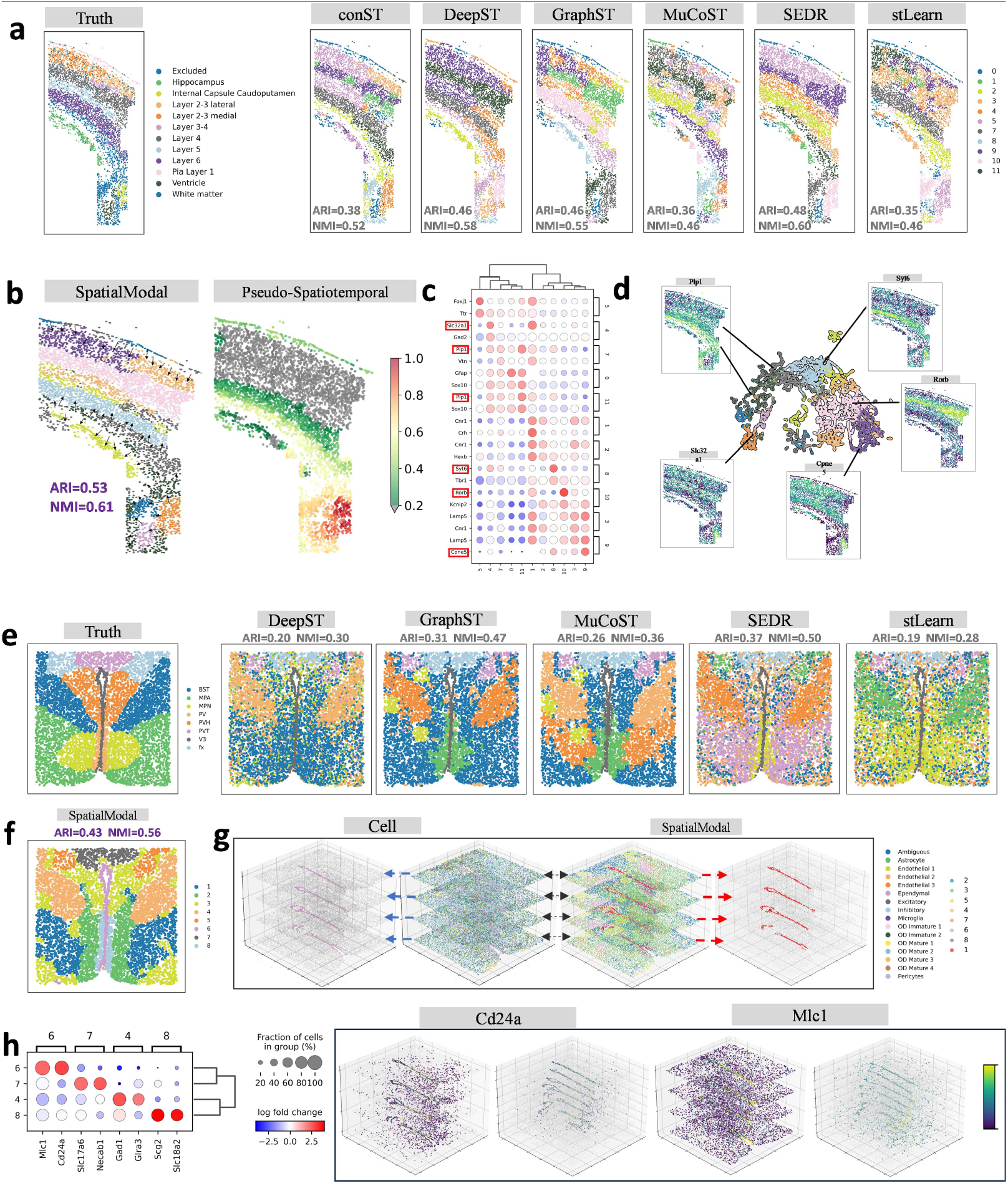
Performance on single-cell-resolution datasets. a. osmFISH somatosensory cortex ground truth and clustering results for six baselines. b. SpatialModal clustering and pseudotemporal trajectories for osmFISH data. c. Bubble plot of domain-specific DEGs. d. UMAP visualization (colored by ground truth) and reconstructed marker gene expression. e. MERFISH hypothalamic preoptic area annotations (Bregma − 0.14) and baseline clustering results. f. SpatialModal clustering for Bregma − 0.14. g. 3D reconstruction of ependymal cells across four Bregma slices and multi-slice clustering results. h. DEGs in domains 4, 6, 7, 8 and 3D expression patterns of *Cd24a* and *Mlc1*. expression reconstruction accuracy, reflected in the highest median Pearson correlation coefficient (Fig. 6e).

We further validate SpatialModal on MERFISH-based^31^ mouse hypothalamic preoptic area data. For Bregma − 0.14 sections, SpatialModal achieves optimal clustering (ARI=0.43, NMI=0.56) (Fig. 5e, 5f), producing anatomically coherent boundaries for V3, PVT, and PV nuclei compared to fragmented domains from DeepST, SEDR, and stLearn. Multi-slice 3D reconstruction (Fig. 5g) reveals domain 1 overlapping precisely with ependymal regions. Expression reconstruction analysis (Fig. 5h) shows noise-reduced spatial patterns for *Cd24a* and *Mlc1*, with enhanced localization to ependymal zones. These results underscore SpatialModal’s capability in resolving single-cell-resolved spatial architectures and cross-slice molecular dynamics.

### VC and Embryo Datasets Analysis with STARmap and Stereo-seq

We apply SpatialModal to the mouse visual cortex dataset generated by the STARmap^32^ platform to evaluate its performance in spatial transcriptomic analysis. Clustering results demonstrate the superiority of SpatialModal over seven benchmark methods in spatial domain delineation (Fig. 6a, Supplementary Fig. S7A). Specifically, SpatialModal-identified domains 0, 6, and 5 exhibit high concordance with manually annotated CC, L6, and L5, respectively. In contrast, stLearn and DeepST merge L5 and L6 into a single category, while conST and SpaGCN fail to resolve clear anatomical boundaries. Quantitative evaluations confirm SpatialModal’s leading performance, achieving the highest scores in ARI (0.58) and NMI (0.7) (Fig. 6a). The dataset’s vertical laminar architecture provides an ideal framework to assess preservation of native spatial organization. We quantify pairwise distance correlations between latent and original spatial coordinates, revealing complete loss of spatial information in conST and suboptimal correlations in GraphST, MuCoST, and stLearn (dispersed density plots deviating from diagonal alignment). DeepST and SEDR show improved yet inferior correlations compared to SpatialModal, which achieves a Pearson correlation coefficient (PCC) of 0.6 (Fig. 6c, 6d). Differential expression analysis of domains 0, 5, and 6 identifies *Bcl6, Rab3c*, and *Nrgn* as representative marker genes (Fig. 6b, Supplementary Fig. S7B). SpatialModal enhances *Bcl6* expression specificity in domain 0 while suppressing its off-target signals, maintains *Rab3c* fidelity in domain 5 with reduced lateral expression artifacts, and extends *Nrgn* localization in domain 6. Comparative analysis confirms SpatialModal’s superior gene

**Figure 6.**
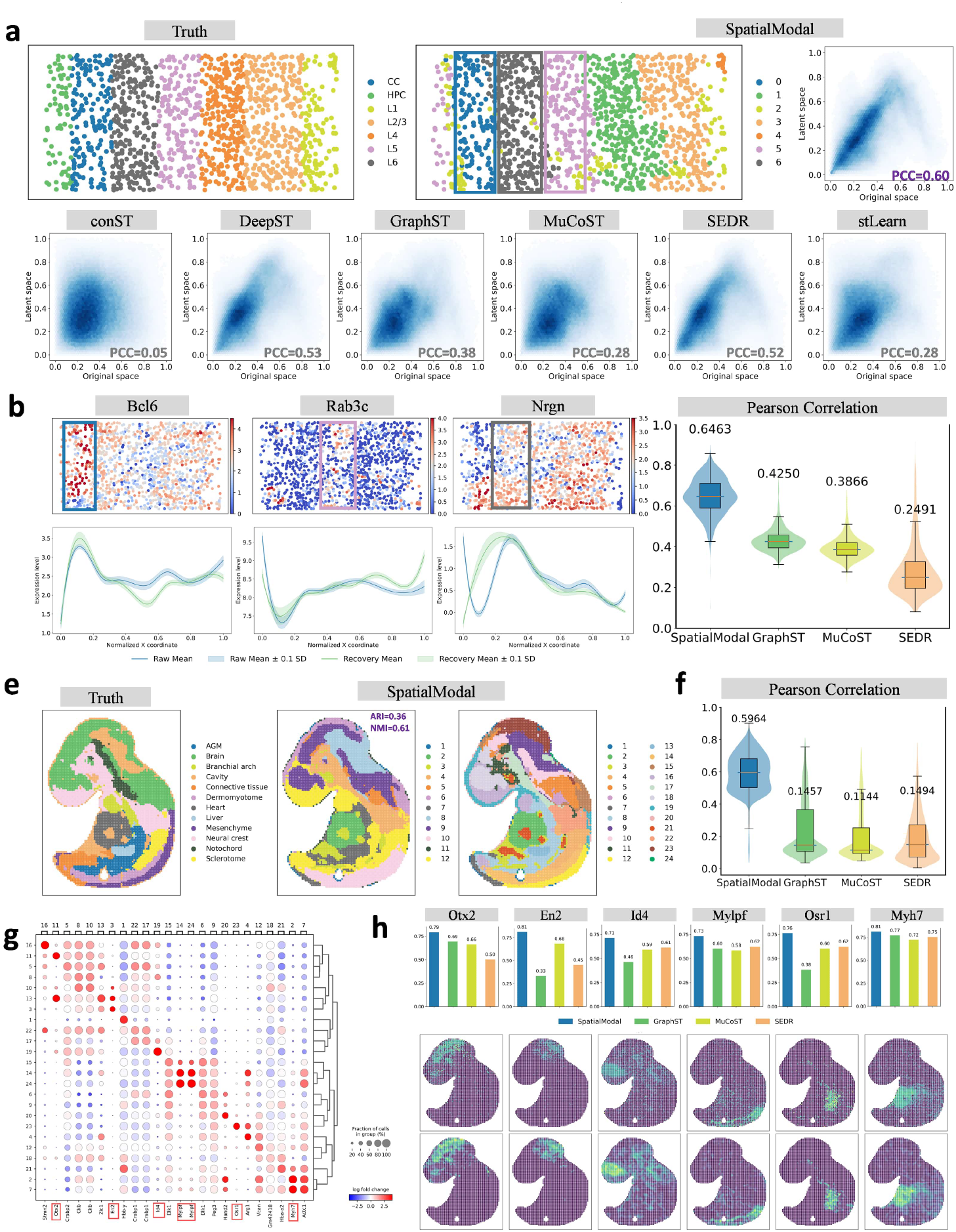
Performance on STARmap and Stereo-seq datasets. a. STARmap visual cortex ground truth, SpatialModal clustering, and density plots of latent-to-spatial correlations. b. Reconstructed *Bcl6, Rab3c*, and *Nrgn* expression profiles along the x-axis; violin plots compare global reconstruction accuracy. c. Stereo-seq embryonic tissue annotations and SpatialModal clustering (*k* = 12, *k* = 24). d. Violin plots of gene reconstruction performance. e. Bubble plot of DEGs across 24 embryonic domains. f. Spatial expression gradients of six marker genes; bar plots show reconstruction accuracy.

We further validate SpatialModal on a Stereo-seq-generated^33^ mouse embryonic dataset^34^. At 12 domains, SpatialModal achieves optimal domain resolution (ARI=0.36, NMI=0.61) (Fig. 6f), outperforming methods like GraphST and SEDR that fail to detect cardiac regions (Fig. 6g). Increasing domain numbers to 24 enables fine-grained annotation alignment, with SpatialModal-identified domains 4 (Sclerotome), 12 (Mesenchyme), 13 (Liver), and 14 (Dermomyotome) matching groundtruth labels (Fig. 6e, Supplementary Fig. S7C). SpatialModal exhibits superior gene expression reconstruction (median Pearson r=0.5964) and precise recovery of spatial expression gradients for differentially expressed genes (Fig. 6h), underscoring its robustness in embryonic tissue analysis.

## Discussion

SpatialModal is an innovative multimodal deep graph learning framework that effectively integrates gene expression data with histological images through hierarchical graph neural networks and contrastive learning, providing fine-grained spatial representations for complex tissue structures. Comprehensive evaluations across diverse datasets highlight its exceptional ability to dissect complex tissue structures with superior precision and adaptability, applicable to various technical platforms. For datasets from the 10x Visium platform with histological images, SpatialModal demonstrates its core strengths. In the DLPFC dataset, it precisely delineates cortical layers, reveals subtle anatomical boundaries, and delivers remarkable detail, offering profound insights into brain tissue organization. In the human breast cancer dataset, SpatialModal uncovers the complex heterogeneity of the tumor microenvironment, accurately mapping specific subregions defined by metabolic and immune features, aligning closely with pathological annotations, and advancing understanding of cancer dynamics. Similarly, in mouse anterior and posterior brain datasets, it adeptly resolves fine structures such as the subiculum and hippocampal regions, providing granular views of molecular and spatial relationships, enriching neurobiological research. These achievements stem from SpatialModals ability to seamlessly integrate molecular and morphological data, enhancing spatial region resolution and reconstructing gene expression with high fidelity. Even without histological images, SpatialModals robust design performs exceptionally, as evidenced by its performance on non-image datasets (osmFISH, MERFISH, STARmap, and Stereo-seq). In the osmFISH dataset of mouse somatosensory cortex, it meticulously distinguishes cortical subregions, capturing complex layer-specific patterns. In the MERFISH dataset of mouse hypothalamic preoptic area, SpatialModal defines critical brain regions with clear, biologically meaningful boundaries. For the STARmap dataset of mouse visual cortex, it accurately reconstructs layered structures, preserving the spatial integrity of functional domains. In the Stereo-seq dataset of mouse embryos, SpatialModal precisely identifies key developmental structures like the heart and liver, facilitating insights into embryonic pattern formation. These results reflect SpatialModals inherent ability to leverage its graph-based architecture and contrastive learning to extract and integrate spatial and molecular signals, even in single-modality settings. By consistently excelling in datasets with and without images, SpatialModal proves its versatility and robustness.

Despite the encouraging performance of our model, certain limitations warrant further discussion. First, our image-based experiments are currently limited to 10x Visium datasets, as this is the only platform providing both histological images and precise pixel-level spatial coordinates for measurement points. This constraint hinders the generalization of our approach to other datasets lacking such imaging information. Second, our current method relies on ResNet152, a deep convolutional neural network pretrained on natural images, to extract histological image features. Although effective, this pretrained model may not fully capture domain-specific patterns inherent to histological images, limiting the biological relevance of learned features. Finally, the complexity of our model architectureintegrating multiple modalities and graph-based learning componentsresults in relatively long computation times during training and inference, potentially restricting scalability to larger datasets. To address these limitations, future research can explore incorporating approximate image reconstruction or registration strategies to include other ST platforms, broadening applicability. Additionally, replacing the current image encoder with domain-adaptive or self-supervised visual models trained directly on histological data may yield more representative image features. To reduce computational burden, we also plan to investigate model simplification strategies, such as knowledge distillation, sparse graph construction, or modular training schemes, to enhance efficiency while maintaining performance.

## Methods

### Overview

SpatialModal is a multi-modal deep graph learning framework designed to achieve higher precision in spatial transcriptomics analysis by integrating gene expression data, spatial coordinates, and histology images (when available). Below, we describe the data preprocessing, model architecture, training strategies, and evaluation protocols in detail.

### Data Preprocessing

To ensure high-quality input data and reduce noise interference, we systematically preprocess gene expression data to filter noise and highlight biological signals. Using the SCANPY package^35^, genes are filtered to retain only those expressed in at least 50 spatial spots with a total expression count ≥ 10, thereby removing low-expression or unreliable genes. Subsequently, 3,000 highly variable genes (HVGs) are selected using the Seurat v3 method, focusing on genes contributing most to tissue heterogeneity. The resulting expression matrix undergoes standardization, log-transformation, and scaling to generate matrix *X* ∈ ℝ^*N×G*^, where *N* is the number of spots and *G* is the number of genes.

For datasets containing histology images (e.g., 10x Visium), we extract image features around each spatial spot. An 80 × 80pixel image patch centered on the spot coordinates is cropped and resized to 299 × 299 pixels to meet the input requirements of the pretrained ResNet-152^36^ model. Deep features are extracted via ResNet-152 to generate an image feature matrix *I* ∈ ℝ^*N×S*^, where *S* is the feature dimension.

To model spatial relationships between spots, a k-nearest neighbor (KNN) graph is constructed based on Euclidean distances between coordinates, reflecting local topological structures in the tissue. Each spot is connected to its *K* nearest neighbors, forming an adjacency matrix *A* ∈ {0, 1}^*N×N*^, where *A*_*i j*_ = 1 if spot *j* is a neighbor of spot *i*, and 0 otherwise. To ensure stability in graph convolution operations, the normalized adjacency matrix is computed as:

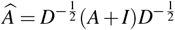

where *D* is the degree matrix and *I* is the identity matrix.

### Data augmentation

To promote cross-modal alignment between gene expression and image features while enhancing model robustness to data variation, we first project gene expression features *X* and image features *I* into a unified dimensional space through two linear mappings. A shared multilayer perceptron (MLP) projection head is then applied to obtain aligned features *Z*_*x*_ and *Z*_*I*_. Using a modality masking strategy, a random subset of spots (default ratio *r* = 0.3) is selected as the mask set ℳ, and their gene expression features are replaced with corresponding image features to generate the augmented feature matrix:

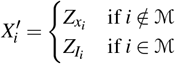

### Hierarchical Encoding

To balance the unique information of gene expression and image modalities as well as their shared features, we designed a hierarchical encoding framework to generate robust joint representations. The enhanced gene expression features *X*^*′*^ and image-aligned features *Z*_*I*_ are first processed through a shared graph convolutional encoder *f*_shared_(·) to capture the interaction between modalities, and then separately passed through modality-specific graph convolutions to extract unique features:

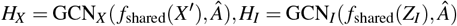

The combined representations *H*_*X*_ and *H*_*I*_ are then fed into a joint encoder to learn the latent distribution. The VGAE encoder calculates the mean µ and variance σ ^2^ :

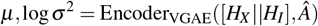

The latent representation *Z* is sampled using reparameterization:

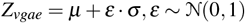

The VGAE decoder reconstructs the graph structure through inner product:

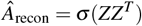

The VGAE loss combines the binary cross-entropy for graph reconstruction and the KL divergence for latent distribution regularization:

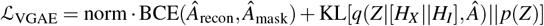

Where BCE(·,·) is the binary cross-entropy loss, 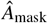 is the masked adjacency matrix, norm is the normalization factor, KL[·||·] denotes the KL divergence, and *p*(*Z*) represents the standard normal prior. After obtaining the combined representation *Z*_fusion_ = [*Z*_vgae_ || *Z*_*i*_], and through linear layer mapping as the final latent representation *Z*_latent_, *Z*_latent_ and *H*_*X*_ are respectively reconstructed through shared linear decoders to obtain gene expression features:

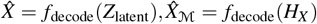

The total reconstruction loss includes the reconstruction errors of all spots and the masked spots:

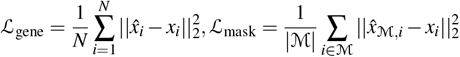

Where *x*_*i*_ and 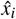 are the original and reconstructed gene expressions of the spots, respectively.

### Topology-aware Contrastive Learning

To enhance the representation’s ability to distinguish between spatial structures and noise, we introduce a topology-aware contrastive learning model^37^. We perform two types of random augmentation (edge and feature random discarding) on the graph embedding *Z*_*X*_ to obtain two enhanced views. After processing through a shared GCN and projection head, we obtain representations *H*_1_ and *H*_2_. Based on these two representations, we consider the embedding of the same node in two views as a positive sample pair, denoted as (*h*_1*i*_, *h*_2*i*_). The sample pair is classified into two categories: one is the embedding of different nodes in the same view, denoted as (*h*_*ni*_, *h*_*n j*_), where *n* ∈ {1, 2}, *j ≠ i* ; the other is the embedding of different nodes in different views, denoted as (*h*_*ai*_, *h*_*b j*_), where *a, b* ∈ {1, 2}, *a ≠ b, j ≠ i*. We use the InfoNCE loss function to maximize the consistency between positive sample pairs and minimize the consistency between negative sample pairs. Taking node *i* as an example, the loss is calculated as follows:

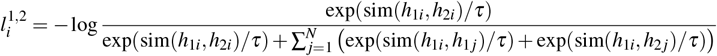

where sim 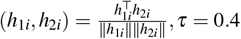 is the temperature parameter, and the denominator is similar. Finally, through these two views, we can obtain the total InfoNCE loss:

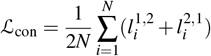

Furthermore, we use the original adjacency matrix 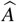 and the gene expression matrix *Z*_*X*_ as inputs. In the next step, we use the GCN as an encoder to train and obtain the low-dimensional embedding *H*_*g*_. Based on this embedding, we use K-means to perform domain prediction for each node and label the regions *L*. Then, we perform feature averaging for all nodes belonging to a domain to obtain the center of each domain *C*. Here, we take the center point and other points in the same domain as positive sample pairs, denoted as 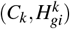, where *k ∈* {1, 2, …, *K*}, *K* represents the number of regions. We take the center point and points in other regions as negative sample pairs, denoted as 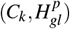, where *k, p* ∈ {1, 2, …, *K*}, *k ≠ p*. Finally, the loss is calculated as follows:

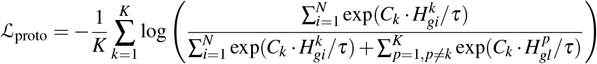

### Total Loss Function

This data representation learning is trained by minimizing the total loss, which includes VGAE loss and contrastive loss. To make the model focus more on gene expression during the training phase, we adopt a two-stage training strategy. The overall training loss of the module is defined as:

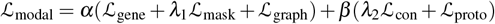

where the default values are λ_1_ = 10, λ_2_ = 0.01. The total number of training epochs is 1000, α = 1, β = 0 when the training epoch is less than 400, and α = 0.5, β = 1 when the training epoch is greater than or equal to 400.

### Multi-Slice Integration Strategy

SpatialModal supports the integration and training of single-slice and multi-slice spatial transcriptomics data. For multiple slices from the same tissue, since there is a lack of direct spatial association between the slices, they need to be integrated to form a unified input. Let *A*_*k*_, *X*_*p*_, *I*_*k*_ represent the adjacency matrix, gene expression matrix, and image feature matrix of slice *k*, respectively. We construct block diagonal matrices *X*_inte_ and *I*_inte_ to store the combined gene expression matrix and the combined image feature matrix, respectively. Specifically:

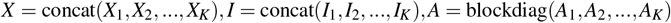

where *K* is the number of slices. The integrated *X, I*, and *A* are used as the input for SpatialModal training.

### Spatial Mapping Based Gene Denoising Module

To improve the biological consistency of denoised gene expression data, inspired by the work of STAGE^38^, we designed a spatial mapping-based denoising module. This module consists of an MLP encoder and a decoder, using batch normalization and ReLU activation. The input is the latent feature *Z*_latent_ ; the encoder outputs the intermediate feature *Z*_mid_ ; the decoder reconstructs the gene expression *X*_rec_. Based on the spatial position of the spots, spatial mapping constraints are established. For a single slice, *Z*_mid_ is 2-dimensional, and the loss is defined as:

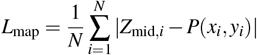

where *x, y* represent the horizontal and vertical coordinates of the spatial points, respectively. Considering the significant differences in the spatial coordinate ranges generated by different platforms for spatial transcriptomics data, we perform a unified normalization operation on the input coordinates, *P* is the normalization operation. For multiple slices, *Z*_mid_ is 3-dimensional,

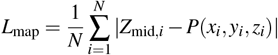

where *z* ∈ [1, 2, 3, …, *M*] represents the slice on which the spatial point is located. The second loss function is the reconstruction loss:

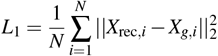

where *X*_*g*_ = [*X*_*g*,1_, *X*_*g*,2_, …, *X*_*g,M*_] is the collection of raw gene expression features from *M* slices. The training uses the Adam optimizer with a learning rate of 0.001, and the learning rate is adjusted using a learning rate scheduler (step size 500, decay rate 1). The overall training loss is defined as:

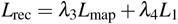

### Method Comparison

#### conST

conST is a multimodal contrastive learning framework for spatial transcriptomic data analysis. It integrates gene expression, spatial coordinates, and histological morphology to learn low-dimensional embeddings, employing contrastive learning to optimize embeddings by maximizing mutual information across local, global, and contextual levels. Parameters used in this study: *k* = 20, *epochs* = 200. For datasets without histology images: *use*_*img* = *False*.

#### DeepST

DeepST is a deep learning-based framework that extracts morphological features from histopathological images using a pretrained neural network. These features are then combined with gene expression and spatial data to generate enhanced transcriptomic profiles. Parameters: *pre*_*epochs* = 500, *epochs* = 500. For datasets lacking morphological data: *use*_*morphological* = *False*.

#### GraphST

GraphST utilizes graph neural networks and self-supervised contrastive learning to learn low-dimensional embeddings by integrating gene expression and spatial adjacency information. Parameters: *k* = 5, *epochs* = 1000.

#### MuCoST

MuCoST employs multi-view graph contrastive learning to fuse gene expression correlations and spatial proximity, enhancing non-local spatial co-expression dependencies. Parameters: *k* = 6, *epochs* = 1000, *corr* = 0.5, *lr* = 0.001.

#### SEDR

SEDR combines deep autoencoders with masked self-supervision to derive low-dimensional gene expression representations, while embedding spatial information through a variational graph autoencoder. Parameters: *k* = 10, *epochs* = 200, *p*_*drop* = 0.2.

#### SpaGCN

SpaGCN constructs weighted graphs integrating gene expression, spatial coordinates, and histological features, followed by graph convolutional aggregation of neighboring spot information.

#### stLearn

stLearn harmonizes gene expression, spatial localization, and histological morphology. It applies spatial smoothing and morphology-aware expression adjustment before performing graph-based clustering for cell-type identification. Parameters: *n*_*comps* = 15, *weights* =^*′*^ *physical*_*distance*^*′*^.

### Evaluation Metrics

#### Adjusted Rand Index(ARI)

The ARI measures similarity between clustering outcomes and ground-truth labels, accounting for chance agreement. It is defined as:

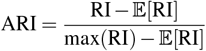

where 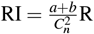 Rand Index is the raw Rand Index, with *a* being the number of sample pairs correctly clustered together, *b* the number correctly separated, and 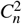 the total sample count. The ARI ranges from [*−*1, 1], where higher values indicate better clustering.

#### Normalized Mutual Information(NMI)

The NMI quantifies consistency between clustering results and true labels via normalized mutual information:

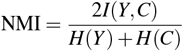

where *I*(*Y,C*) is mutual information between true labels *Y* and domains *C*, and *H*(*Y*), *H*(*C*) denote their respective entropies. Values range from [0, 1], with higher values reflecting superior agreement.

#### Homogeneity

This metric assesses whether each cluster contains samples from a single class:

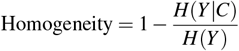

where *H*(*Y*|*C*) is the conditional entropy of labels *Y* given domains *C*, and *H*(*Y*) is the label entropy. Values span [0, 1], with higher scores indicating better homogeneity.

#### Purity-score.^24^

Purity evaluates the dominance of majority classes within domains:

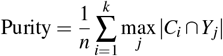

where *n* is the total samples, *k* the number of domains, *C*_*i*_ the *i* -th cluster, and *Y*_*j*_ the *j* -th true class. Values range from [0, 1], with higher scores indicating better class separation.

#### Silhouette score.^39^

This score measures intra-cluster tightness and inter-cluster separation per sample:

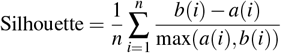

where *a*(*i*) is the mean distance between sample *i* and others in its cluster, and *b*(*i*) the smallest mean distance to samples in other domains. The global score is the average over all samples, ranging from [−1, 1], where higher values denote better clustering.

#### Davies-Bouldin Score.^40^

This metric evaluates the ratio of intra-cluster compactness to inter-cluster separation:

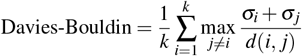

where σ_*i*_ is the average distance of cluster *i* ‘s samples to its centroid, and *d*(*i, j*) the distance between centroids of domains *i* and *j*. Lower values indicate better clustering, with the ideal score being 0.

#### Pearson Correlation Coefficient(PCC)

The PCC quantifies linear correlation between variables:

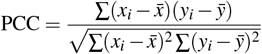

where *x*_*i*_ and *y*_*i*_ are paired observations, and 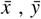 their means. Values range from [− 1, 1], with *r >* 0 indicating positive correlation and higher absolute values representing stronger associations.

### Downstream analysis methods

#### K-means clustering

K-means clustering is a distance-based algorithm that partitions data points into *K* domains by minimizing the within-cluster sum of squared distances to cluster centroids. Using the scikit-learn implementation, we apply K-means (random seed=0) to latent embeddings for initial clustering. To enhance spatial coherence, a `refine_label` function adjusts cluster assignments by majority voting among 50 nearest neighbors in Euclidean space.

Datasets applied: Human DLPFC, breast cancer, mouse somatosensory cortex, and mouse visual cortex datasets.

#### Model-based clustering (Mclust)

Model-based clustering assumes data generation from a Gaussian mixture model (GMM), where parameters are estimated via expectation-maximization (EM) to assign cluster labels. Using the mclust R package (model=“EEE”, random seed=2025), we enforce ellipsoidal clusters with equal covariance structures.

Datasets applied: Mouse anterior/posterior brain, hypothalamic preoptic area, and embryonic datasets.

#### Uniform manifold approximation and projection (UMAP)

UMAP is a nonlinear dimensionality reduction technique that preserves local and global topological structures by modeling high-dimensional data as a weighted graph. We apply UMAP for 2D visualization of latent embeddings to elucidate spatial domain relationships.

#### Partition-based graph abstraction (PAGA)

PAGA constructs simplified graphs to quantify connectivity between annotated clusters, enabling trajectory inference by resolving transitional states and lineage relationships.

#### Diffusion pseudotime (DPT)

DPT computes pseudotemporal ordering using diffusion operators derived from cell similarity graphs. Starting from user-defined root cells, cumulative diffusion distances are calculated via random walks, implemented using *scanpy*.*tl*.*di f f map* and *scanpy*.*tl*.*dpt*.

#### Ripleys L function

Ripleys L-function evaluates spatial clustering or dispersion of point patterns. The L-function is defined as:

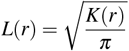

where *K*(*r*) estimates the average number of neighboring points within radius *r*. Values *L*(*r*) *> r* indicate clustering, *L*(*r*) *< r* indicates dispersion, and *L*(*r*) ≈ *r* indicates randomness.

#### CellChat

CellChat infers cell-cell communication networks by analyzing ligand-receptor interactions from the curated CellChatDB. It quantifies communication probabilities using mass action kinetics, integrates co-factors/coreceptor regulation, and visualizes signaling dynamics.

#### Identification of differentially expressed genes

We identify spatial domain-specific DEGs using *scanpy*.*tl*.*rank*_*genes*_*groups* with the Wilcoxon rank-sum test. Genes are ranked by absolute z-scores (|*z*| *>* 3), adjusted p-values (Benjamini-Hochberg FDR *<* 10^*−*3^), and log-fold changes (| log *FC*| *>* 1).

#### GO enrichment analysis

GO enrichment analysis detects statistically overrepresented biological processes, molecular functions, and cellular components among DEGs using hypergeometric testing. Significantly enriched terms (FDR-adjusted p *<* 0.05) are reported.

## Datasets

### Human prefrontal cortex data by 10x Visium

The DLPFC dataset, generated using the 10x Visium platform, was created by researchers at the Lieber Institute for Brain Development (LIBD). It comprises 12 tissue sections (labeled 151507151510, 151669151672, and 151673151676) from three healthy adult donors, capturing 4,226, 4,384, 4,789, 4,634, 3,661, 3,498, 4,110, 4,015, 3,639, 3,673, 3,592, and 3,460 spatial spots, respectively. The first and third section groups cover six cortical layers (Layer 16) and white matter (WM), while the second group includes Layers 36 and WM. Gene expression data were captured via barcoded spatial arrays, with manually annotated spatial domains serving as ground truth for benchmarking spatial clustering accuracy. Data availability: https://github.com/LieberInstitute/spatialDLPFC.

### Human breast cancer data by 10x Visium

This 10x Visium-generated dataset includes 3,798 spatial spots from human breast cancer tissues, with pathological annotations categorizing regions into ductal/lobular carcinoma in situ (DCIS/LCIS), invasive ductal carcinoma (IDC), healthy tissue, and tumor margins. It enables investigations into molecular transitions from normal to malignant states and identification of therapeutic targets. Data availability: https://support.10xgenomics.com/spatial-gene-expression/datasets/1.1.0.

### Mouse brain anterior and posterior data by 10x Visium

The mouse anterior and posterior brain datasets, generated via 10x Visium, consist of 3,289 and 2,825 spatial spots, respectively. The anterior dataset includes manual annotations for the hypothalamus, hippocampus, and other subcortical structures, enabling validation of methods for resolving complex neuroanatomical architectures. Data availability: https://www.10xgenomics.com/resources/datasets.

### Mouse somatosensory cortex data by osmFISH

The osmFISH dataset provides single-cell-resolution spatial transcriptomic profiles for 4,839 cells across 33 genes in the mouse somatosensory cortex. Manually annotated layers include Pia Layer 1, Layer 2-3 (medial/lateral), Layer 4, Layer 5, Layer 6, hippocampus, ventricle, and white matter. This layered annotation supports studies of cortical gene expression gradients and cellular spatial organization. Data availability: http://linnarssonlab.org/osmFISH/.

### Mouse hypothalamic preoptic data by MERFISH

This MERFISH dataset contains single-cell spatial transcriptomes of the mouse hypothalamic preoptic area across four continuous coronal sections (Bregma coordinates: -0.04, -0.09, -0.14, -0.19), with 6,154, 6,185, 5,926, and 6,507 cells per slice. The -0.14 Bregma section includes fine-grained annotations for eight functional regions: BST, MPA, MPN, PV, PVH, PVT, V3, and fx. Data availability: https://datadryad.org/dataset/doi:10.5061/dryad.8t8s248.

### Mouse visual cortex data by STARmap

The STARmap-generated dataset profiles 1,207 spatial spots and 1,020 genes in the mouse primary visual cortex (V1), with manual annotations for seven cortical layers (L1, L2/3, L4, L5, L6), corpus callosum (CC), and hippocampal cortex (HPC). This layered architecture supports studies of neuronal gene regulation and spatial functional specialization. Data availability: https://kangaroo-goby.squarespace.com/data.

### Mouse embryo data by Stereo-seq

The Stereo-seq mouse embryonic dataset contains 5,319 spatial spots and thousands of genes across multiple embryonic tissues. Manual annotations delineate 12 spatial domains, including heart, brain, liver, dermomyotome, and sclerotome, enabling developmental gene expression studies. Data availability: https://db.cngb.org/stomics/mosta/.

## Acknowledgements

This work is supported in part by the Nation Natural Science Foundation of China[62433016, 62202383], and the National Key Research and Development Program of China [2022YFD1801200].

## Author contributions

D.Z. (Dongmin Zhao) developed the SpatialModal framework, conducted all experiments, and wrote the manuscript. X.L. (Xingyi Li) supervised the entire project, guided model design and experiments, and provided critical revisions to the manuscript. J.Z. (Junnan Zhu) provided detailed feedback and revisions on the manuscript. M.L. (Min Li) provided insightful suggestions on the experimental design and result interpretation. F.W. (Feng Wei) provided supporting datasets and contributed to the experimental setup. Y.Q. (Yang Qi), Y.C. (Yiqi Chen), and Y.W. (Yingfu Wu) contributed to the literature review and provided valuable feedback during manuscript preparation. X.J. (Xiangting Jia), G.D. (Gaoyuan Du), and J.X. (Jialuo Xu) assisted with model validation, technical discussions, and resolving implementation issues. X.S. (Xuequn Shang) provided overall academic support and institutional resources. All authors read and approved the final manuscript.

