## Supplementary Figures and Tables for "A Multimodal Graph Learning Framework for Versatile Spatial Transcriptomics Analysis with SpatialModal"

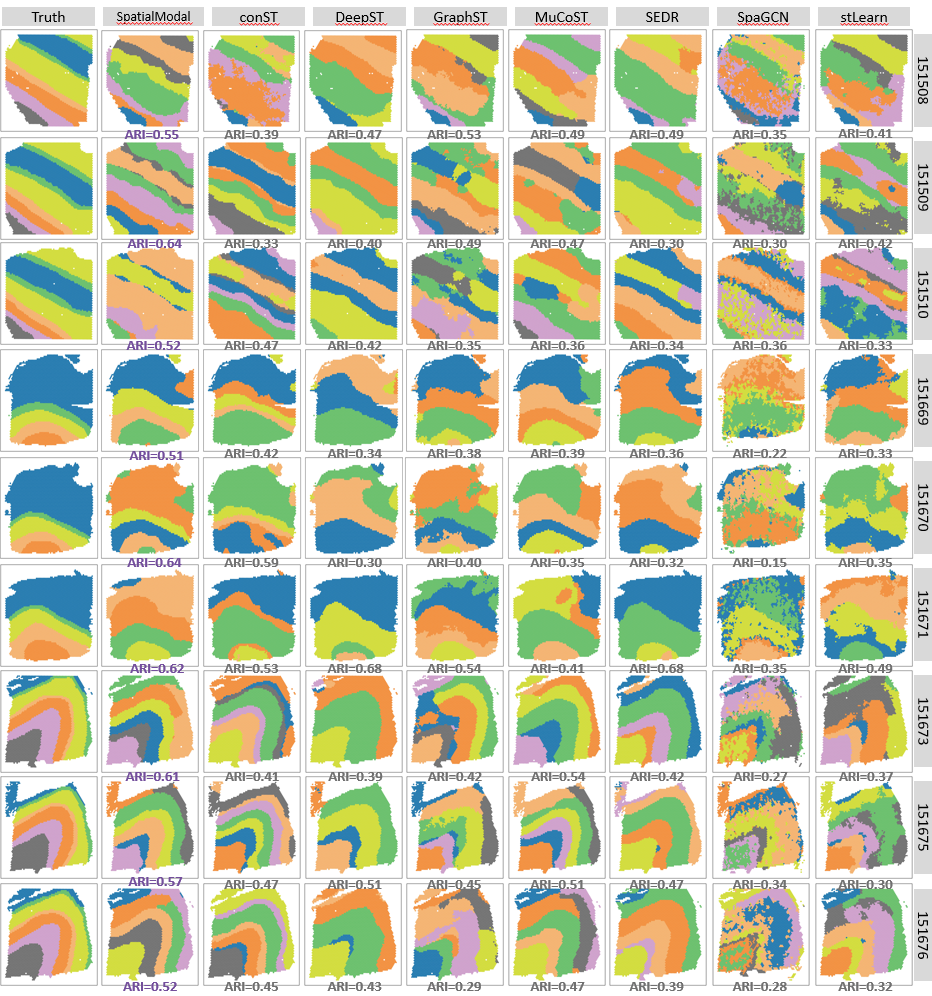

**Figure S1:**Ground-truth annotations and spatial clustering results for nine additional DLPFC slices, comparing SpatialModal with seven baseline methods.

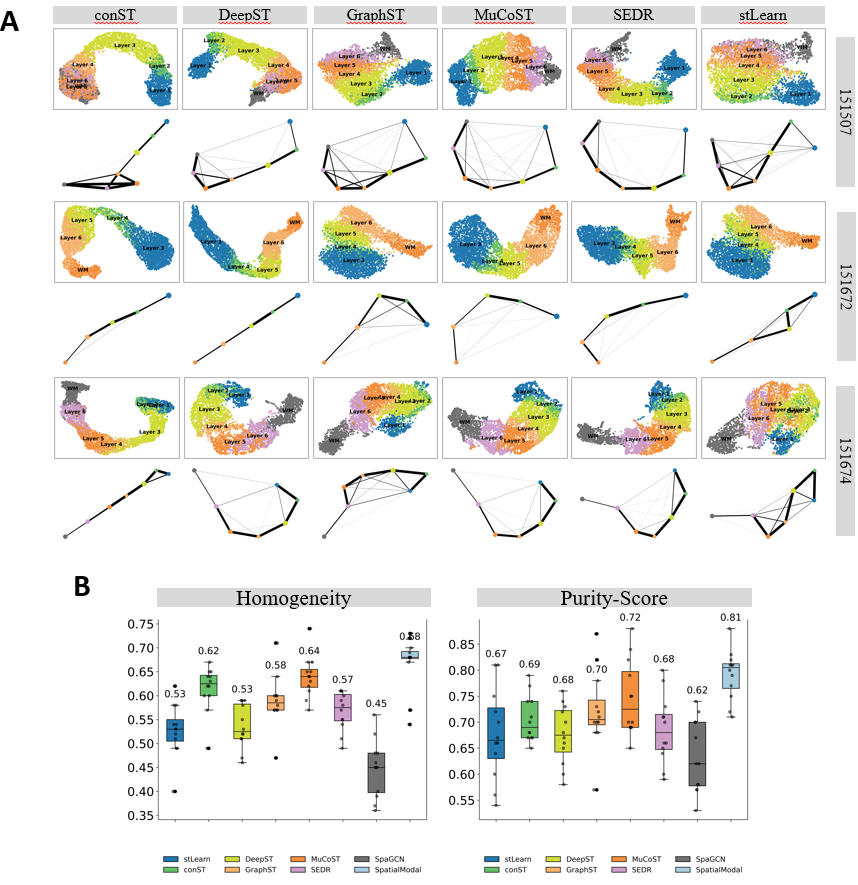

**Figure S2:** A.UMAP embeddings (colored by ground truth) and PAGA trajectories for three DLPFC slices across six baseline methods.B.Boxplots comparing Homogeneity and Purity scores across 12 DLPFC slices for all methods.

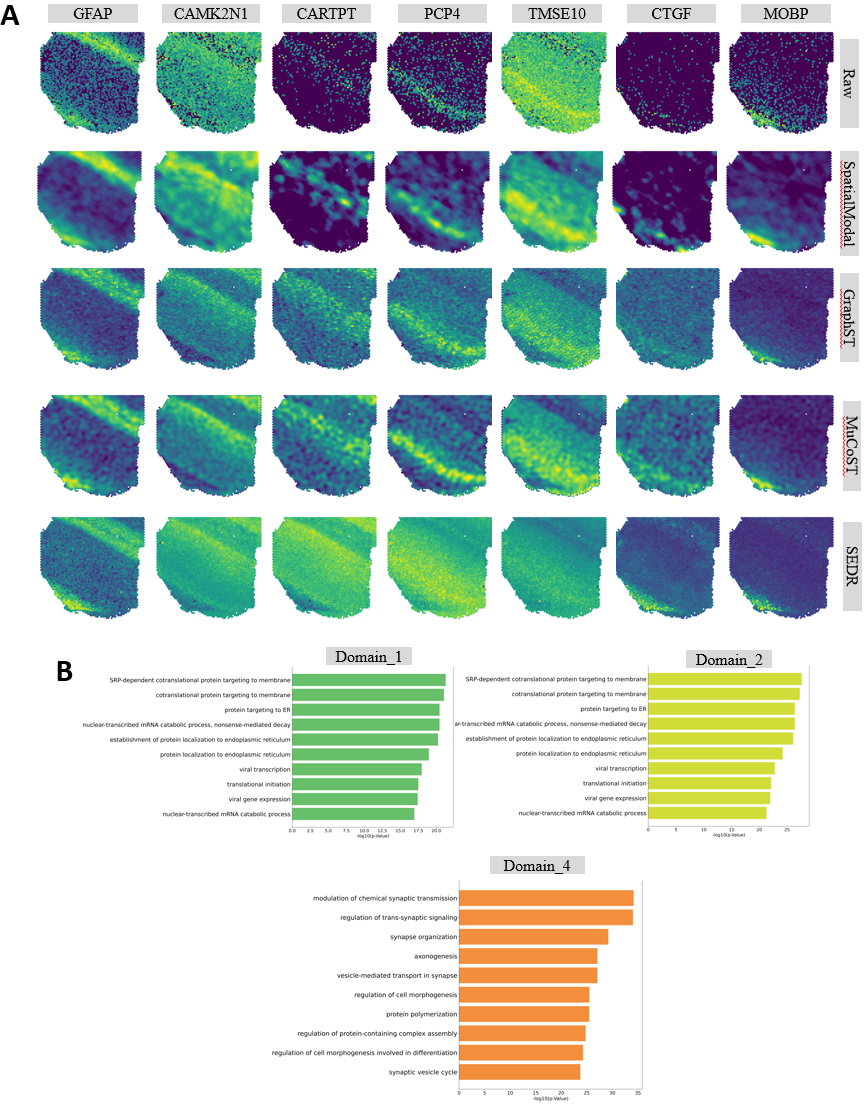

**Figure S3:** A.Spatial expression patterns of seven layer-specific marker genes (raw and reconstructed by SpatialModal and three baselines).B.Top 10 enriched GO terms for clusters 1, 2, and 4 identified in human DLPFC data.

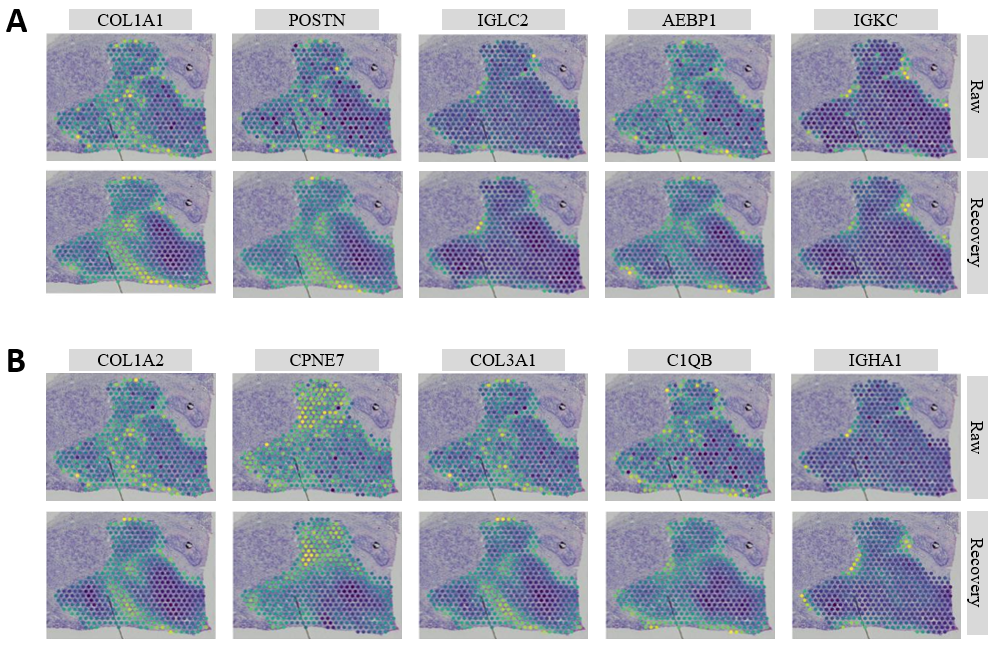

**Figure S4:** Spatial expression patterns of top five upregulated genes in cluster 0 versus cluster 8 (raw and reconstructed). B.Spatial expression patterns of top five upregulated genes in cluster 8 versus cluster 0 (raw and reconstructed).

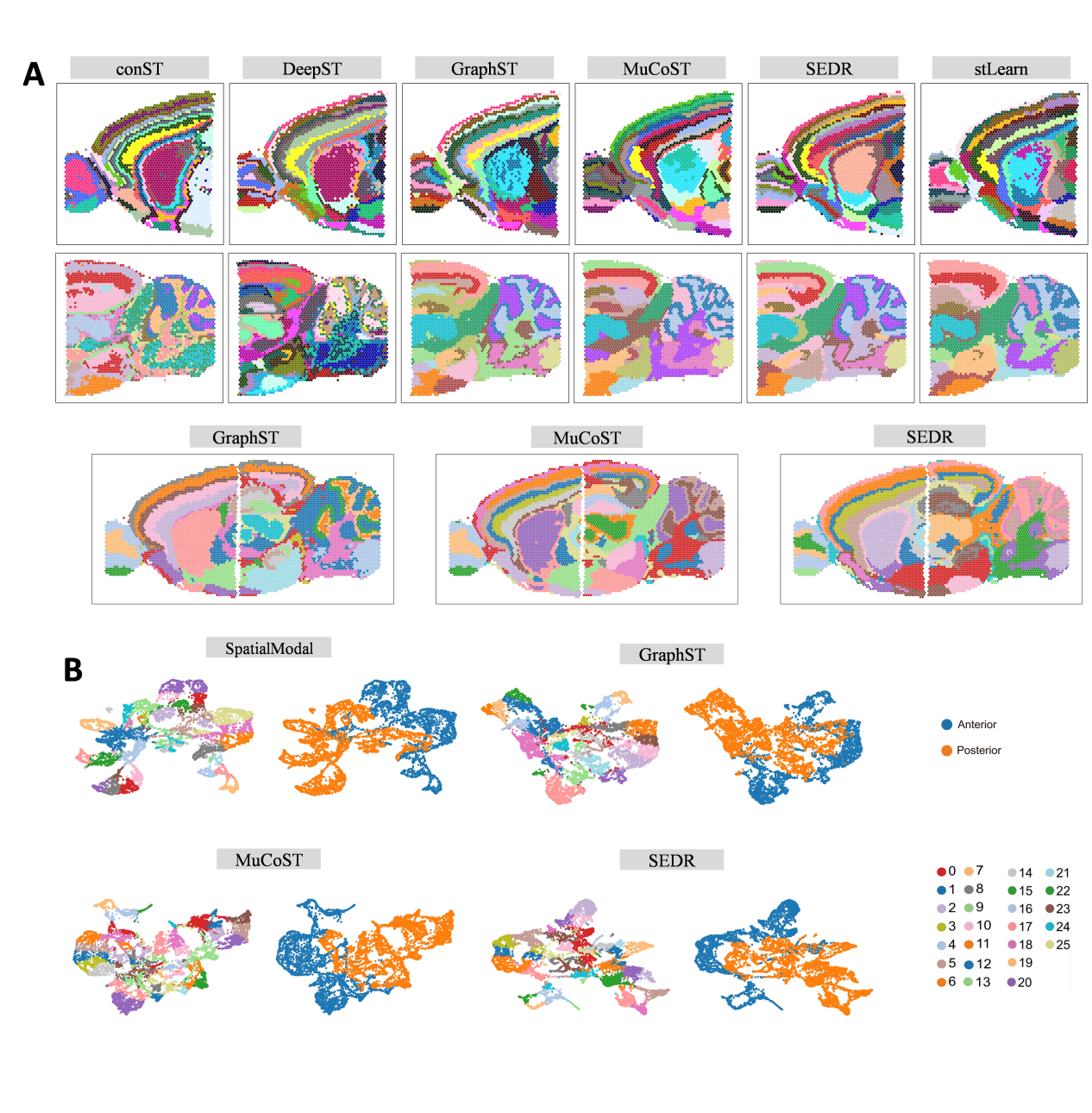

**Figure S5:** A.Spatial clustering results for six baselines on individual and combined mouse forebrain/hindbrain datasets. B.UMAP visualizations of latent embeddings from cross-slice integration, colored by anatomical origin (forebrain/hindbrain) and derived spatial domains.

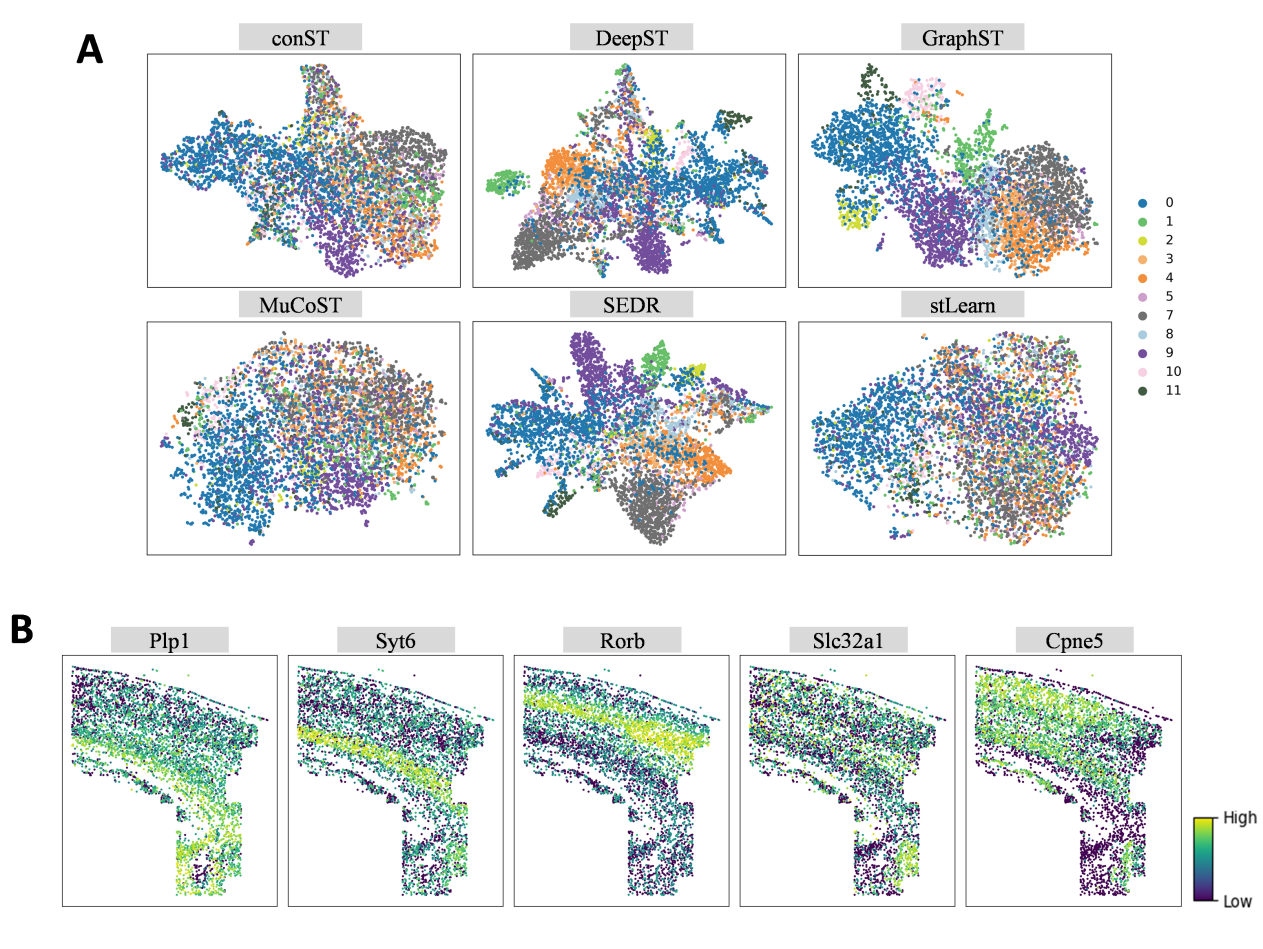

**Figure S6:** A.UMAP embeddings (colored by inferred domains) for six baselines on the mouse somatosensory cortex dataset.B.Spatial expression gradients of five differentially expressed genes (raw data).

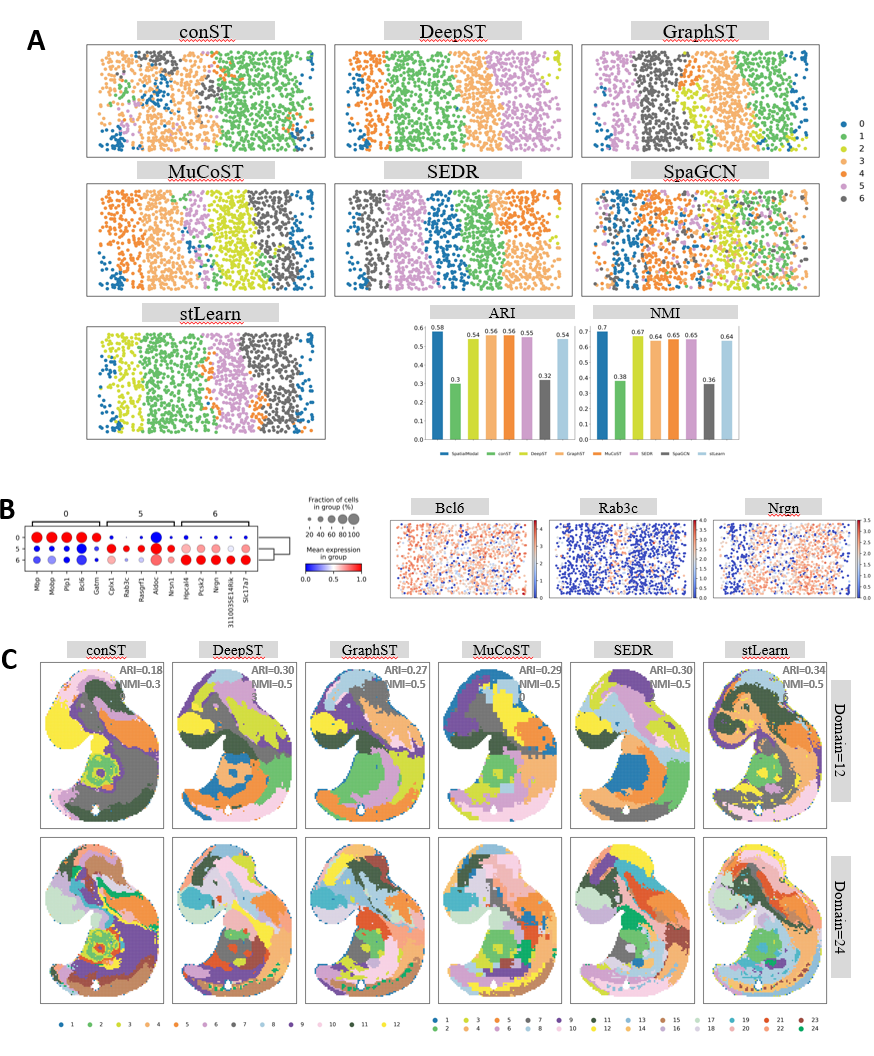

Figure S7: A.Spatial clustering results for seven baselines on the STARmap visual cortex dataset, with ARI/NMI comparisons.B.Bubble plot of DEGs in STARmap clusters 0, 5, and 6, alongside raw expression patterns for three markers.C.Spatial clustering results for six baselines on the Stereo-seq embryonic dataset (k=12, k=24).

**Supplementary Tables**

Table S1：Normalized Mutual Information (NMI) values for SpatialModal and seven baselines across 12 DLPFC slices.

|  | **SpatialModal** | **stLearn** | **conST** | **DeepST** | **GraphST** | **MuCoST** | **SEDR** | **SpaGCN** |
| --- | --- | --- | --- | --- | --- | --- | --- | --- |
| **151507** | 0.687 | 0.541 | 0.52 | 0.606 | 0.653 | 0.631 | 0.664 | 0.456 |
| **151508** | 0.59 | 0.53 | 0.521 | 0.641 | 0.583 | 0.606 | 0.549 | 0.459 |
| **151509** | 0.666 | 0.557 | 0.62 | 0.587 | 0.534 | 0.582 | 0.616 | 0.533 |
| **151510** | 0.624 | 0.509 | 0.581 | 0.564 | 0.528 | 0.542 | 0.629 | 0.471 |
| **151669** | 0.623 | 0.474 | 0.554 | 0.576 | 0.543 | 0.581 | 0.618 | 0.359 |
| **151670** | 0.623 | 0.488 | 0.537 | 0.487 | 0.496 | 0.556 | 0.504 | 0.301 |
| **151671** | 0.685 | 0.588 | 0.641 | 0.655 | 0.673 | 0.612 | 0.619 | 0.458 |
| **151672** | 0.756 | 0.554 | 0.669 | 0.665 | 0.598 | 0.711 | 0.587 | 0.374 |
| **151673** | 0.663 | 0.57 | 0.616 | 0.569 | 0.587 | 0.633 | 0.64 | 0.454 |
| **151674** | 0.692 | 0.402 | 0.63 | 0.621 | 0.559 | 0.627 | 0.69 | 0.369 |
| **151675** | 0.696 | 0.496 | 0.61 | 0.646 | 0.596 | 0.659 | 0.659 | 0.564 |
| **151676** | 0.607 | 0.507 | 0.631 | 0.669 | 0.468 | 0.599 | 0.687 | 0.443 |

Table S2：Homogeneity metrics for all methods across 12 DLPFC slices.

|  | **SpatialModal** | **stLearn** | **conST** | **DeepST** | **GraphST** | **MuCoST** | **SEDR** | **SpaGCN** |
| --- | --- | --- | --- | --- | --- | --- | --- | --- |
| **151507** | 0.692 | 0.534 | 0.499 | 0.501 | 0.645 | 0.64 | 0.608 | 0.463 |
| **151508** | 0.613 | 0.536 | 0.466 | 0.592 | 0.58 | 0.617 | 0.492 | 0.462 |
| **151509** | 0.657 | 0.581 | 0.65 | 0.516 | 0.567 | 0.616 | 0.559 | 0.562 |
| **151510** | 0.738 | 0.513 | 0.618 | 0.539 | 0.568 | 0.585 | 0.615 | 0.478 |
| **151669** | 0.575 | 0.523 | 0.624 | 0.476 | 0.593 | 0.654 | 0.55 | 0.397 |
| **151670** | 0.574 | 0.533 | 0.621 | 0.496 | 0.569 | 0.65 | 0.541 | 0.357 |
| **151671** | 0.686 | 0.618 | 0.603 | 0.553 | 0.714 | 0.632 | 0.581 | 0.472 |
| **151672** | 0.784 | 0.543 | 0.581 | 0.573 | 0.601 | 0.738 | 0.546 | 0.387 |
| **151673** | 0.652 | 0.577 | 0.59 | 0.493 | 0.596 | 0.63 | 0.612 | 0.453 |
| **151674** | 0.686 | 0.397 | 0.626 | 0.541 | 0.566 | 0.641 | 0.67 | 0.373 |
| **151675** | 0.698 | 0.488 | 0.618 | 0.565 | 0.6 | 0.669 | 0.627 | 0.564 |
| **151676** | 0.653 | 0.488 | 0.604 | 0.617 | 0.468 | 0.605 | 0.655 | 0.448 |

Table S3：Purity-Score comparisons across methods for 12 DLPFC slices.

|  | **SpatialModal** | **stLearn** | **conST** | **DeepST** | **GraphST** | **MuCoST** | **SEDR** | **SpaGCN** |
| --- | --- | --- | --- | --- | --- | --- | --- | --- |
| **151507** | 0.806 | 0.645 | 0.603 | 0.608 | 0.697 | 0.701 | 0.653 | 0.584 |
| **151508** | 0.743 | 0.666 | 0.61 | 0.683 | 0.704 | 0.699 | 0.596 | 0.588 |
| **151509** | 0.792 | 0.697 | 0.752 | 0.656 | 0.709 | 0.708 | 0.689 | 0.701 |
| **151510** | 0.892 | 0.659 | 0.695 | 0.63 | 0.683 | 0.705 | 0.708 | 0.673 |
| **151669** | 0.752 | 0.752 | 0.789 | 0.726 | 0.777 | 0.818 | 0.773 | 0.723 |
| **151670** | 0.805 | 0.805 | 0.786 | 0.77 | 0.821 | 0.836 | 0.803 | 0.734 |
| **151671** | 0.847 | 0.809 | 0.734 | 0.727 | 0.867 | 0.755 | 0.733 | 0.694 |
| **151672** | 0.895 | 0.725 | 0.705 | 0.721 | 0.733 | 0.881 | 0.682 | 0.62 |
| **151673** | 0.708 | 0.657 | 0.626 | 0.603 | 0.679 | 0.71 | 0.667 | 0.565 |
| 151674 | 0.746 | 0.56 | 0.686 | 0.583 | 0.705 | 0.749 | 0.74 | 0.53 |
| 151675 | 0.782 | 0.539 | 0.668 | 0.673 | 0.724 | 0.691 | 0.645 | 0.628 |
| 151676 | 0.734 | 0.597 | 0.636 | 0.669 | 0.568 | 0.696 | 0.731 | 0.583 |

Table S4：Upregulated differentially expressed genes (DEGs) in cluster 0 versus cluster 8 (human breast cancer dataset).

| **name** | **z-score** | **p-value** | **pvals_adj** | **logfoldchanges** |
| --- | --- | --- | --- | --- |
| COL1A1 | 9.027 | 1.757E-19 | 1.054E-16 | 1.192 |
| POSTN | 8.994 | 2.382E-19 | 1.191E-16 | 1.333 |
| AEBP1 | 8.843 | 9.289E-19 | 2.470E-16 | 1.040 |
| COL1A2 | 8.490 | 2.073E-17 | 3.658E-15 | 1.101 |
| COL3A1 | 8.200 | 2.404E-16 | 3.606E-14 | 1.072 |
| SFRP2 | 7.705 | 1.307E-14 | 1.634E-12 | 1.119 |
| DCN | 7.513 | 5.799E-14 | 5.428E-12 | 1.068 |
| SERPINF1 | 6.603 | 4.030E-11 | 2.325E-09 | 1.059 |
| SLC40A1 | 5.489 | 4.044E-08 | 1.225E-06 | 1.157 |
| MMP1 | 5.430 | 5.640E-08 | 1.612E-06 | 2.146 |
| CCDC80 | 5.378 | 7.514E-08 | 2.107E-06 | 1.234 |
| COL12A1 | 5.164 | 2.412E-07 | 5.981E-06 | 1.012 |
| OLFML3 | 4.850 | 1.233E-06 | 2.535E-05 | 1.269 |
| ADA2 | 4.335 | 1.460E-05 | 2.368E-04 | 1.013 |
| COL11A1 | 4.333 | 1.471E-05 | 2.372E-04 | 1.148 |
| SFRP4 | 4.281 | 1.858E-05 | 2.904E-04 | 1.052 |
| FPR3 | 4.043 | 5.285E-05 | 7.207E-04 | 1.490 |
| TPSB2 | 3.861 | 1.128E-04 | 1.399E-03 | 1.436 |
| CDH11 | 3.843 | 1.215E-04 | 1.493E-03 | 1.057 |
| TGFBR2 | 3.531 | 4.142E-04 | 4.169E-03 | 1.067 |
| FYB1 | 3.483 | 4.952E-04 | 4.919E-03 | 1.529 |
| LRRC15 | 3.443 | 5.744E-04 | 5.523E-03 | 1.150 |
| CD2 | 3.414 | 6.395E-04 | 6.033E-03 | 1.349 |
| TNC | 3.384 | 7.145E-04 | 6.636E-03 | 1.284 |
| COL10A1 | 3.368 | 7.565E-04 | 6.962E-03 | 1.036 |
| JCHAIN | 3.321 | 8.968E-04 | 7.983E-03 | 1.390 |

Table S5：Upregulated differentially expressed genes (DEGs) in cluster 8 versus cluster 0 (human breast cancer dataset).

| **name** | **z-score** | **p-value** | **pvals_adj** | **logfoldchanges** |
| --- | --- | --- | --- | --- |
| COX6C | 10.460 | 1.319E-25 | 3.958E-22 | 1.121 |
| IER3 | 10.307 | 6.534E-25 | 9.800E-22 | 1.088 |
| TFF1 | 9.233 | 2.638E-20 | 2.638E-17 | 1.026 |
| ABCC12 | 9.189 | 3.951E-20 | 2.963E-17 | 2.655 |
| ABCC11 | 8.880 | 6.676E-19 | 2.003E-16 | 1.930 |
| GSTM3 | 8.882 | 6.539E-19 | 2.003E-16 | 1.605 |
| AC093001.1 | 8.629 | 6.195E-18 | 1.327E-15 | 1.529 |
| ACADSB | 8.536 | 1.384E-17 | 2.595E-15 | 1.022 |
| BAG3 | 8.383 | 5.177E-17 | 8.628E-15 | 1.384 |
| PTGES | 7.675 | 1.656E-14 | 1.988E-12 | 1.402 |
| ECM1 | 7.588 | 3.246E-14 | 3.438E-12 | 1.378 |
| ALDH3B2 | 7.223 | 5.103E-13 | 3.734E-11 | 1.264 |
| SIAH2 | 6.664 | 2.672E-11 | 1.636E-09 | 1.031 |
| NAT1 | 6.410 | 1.451E-10 | 7.776E-09 | 1.131 |
| RBM24 | 6.276 | 3.480E-10 | 1.657E-08 | 1.727 |
| C17orf58 | 6.111 | 9.891E-10 | 4.239E-08 | 1.201 |
| APOD | 6.082 | 1.184E-09 | 4.901E-08 | 1.340 |
| PIP | 5.839 | 5.246E-09 | 1.830E-07 | 1.221 |
| MAPT | 5.559 | 2.717E-08 | 8.861E-07 | 1.240 |
| MGST1 | 5.475 | 4.372E-08 | 1.304E-06 | 1.061 |
| UNC13B | 5.428 | 5.714E-08 | 1.617E-06 | 1.283 |
| PLEKHD1 | 5.271 | 1.357E-07 | 3.602E-06 | 1.129 |
| SGK3 | 5.047 | 4.497E-07 | 1.062E-05 | 1.178 |
| DIO1 | 4.938 | 7.876E-07 | 1.712E-05 | 1.089 |
| STC1 | 4.745 | 2.080E-06 | 4.133E-05 | 1.249 |
| SGCG | 4.417 | 9.987E-06 | 1.665E-04 | 1.396 |
| HRASLS2 | 4.360 | 1.301E-05 | 2.121E-04 | 1.278 |
| CCT6B | 4.159 | 3.203E-05 | 4.665E-04 | 1.708 |
| ADAMTS15 | 4.125 | 3.711E-05 | 5.227E-04 | 1.099 |
| REEP1 | 4.048 | 5.165E-05 | 7.076E-04 | 1.587 |
| ABCA12 | 4.011 | 6.041E-05 | 8.164E-04 | 2.287 |
| PLPPR3 | 3.924 | 8.705E-05 | 1.108E-03 | 1.036 |
| Z82246.1 | 3.741 | 1.832E-04 | 2.074E-03 | 1.010 |
| CRISP2 | 3.645 | 2.672E-04 | 2.871E-03 | 1.014 |
| S1PR3 | 3.642 | 2.704E-04 | 2.887E-03 | 1.054 |
| SLC1A2 | 3.399 | 6.775E-04 | 6.331E-03 | 1.038 |
| C1orf115 | 3.296 | 9.817E-04 | 8.463E-03 | 1.093 |
| MESP2 | 3.269 | 1.078E-03 | 9.112E-03 | 1.095 |
